# On the effect of applied lateral tension on the deformation energetics of biological membranes and the lipid dynamics within

**DOI:** 10.64898/2026.03.06.710217

**Authors:** Yein Christina Park, Giacomo Fiorin, José D. Faraldo-Gómez

## Abstract

A broad range of cellular functions involve transient or persistent changes in the morphology of lipid membranes, from the organellar to the molecular scale. By and large, the thermodynamics of protein-mediated remodeling, which often distort the membrane beyond its thermally accessible undulations, remain to be understood. Molecular Dynamics simulations enhanced by advanced sampling methods are uniquely suited to examine and quantitate these phenomena. Here, we leverage the Multi-Map simulation method to directly quantify how applied lateral tension impacts the energetics of both global and localized membrane perturbations induced extrinsically. We obtain free-energy profiles of curvature and thickness deformations with and without tension, which allow us to test the quadratic approximations of Helfrich-based continuum theories. We find that these continuum models are applicable up to ∼ 20 k_B_T, beyond which they under-estimate the free-energy cost of bending. However, we find that the effect of tension on these deformations is primarily driven by changes in the projected area of the membrane, and therefore consistent with continuum theory, even beyond 30 k_B_T. These changes outpace the effects of tension on lipid packing and diffusion, and importantly we find that tension levels preceding rupture (≤ 10 mN/m) do not impede complete lipid mixing. Finally, we investigate the free-energy landscapes of membrane thickness and lipid enrichment with a heterogeneous lipid composition and show that strong enrichment can be sustained even under steady exchange of the constituent lipid molecules.

**Statement of Significance:** Cells have evolved the ability to sense mechanical forces through specialized proteins embedded in their membranes. How exactly the membrane transduces these stimuli to the proteins therein has been unclear. Using state-of-the-art molecular simulations, we quantify the free-energy landscape of curvature and thickness deformations, varying either the applied lateral tension or lipid composition. Our results show that stretching a membrane does not exert an outward force on individual lipid molecules, even when these are biased toward a specific region of the membrane. Instead, lateral tension alters the membrane’s resistance to remodeling by penalizing changes in projected area, suggesting that membrane deformation energetics, and not lipid residence, drives the tension-responsive conformational changes in stretch-activated proteins.

## Introduction

Membrane protein function and regulation are deeply connected to the composition, morphology, and plasticity of the lipid bilayer. Indeed, it is increasingly apparent, based on computer-simulation studies and experimental high-resolution structural data, that the shape and chemical composition of the membrane in the vicinity of proteins are just as diverse as the folds of the proteins themselves. Localized shape deformations, for example, develop to prevent the exposure of polar amino-acids to the membrane interior, and to shield hydrophobic amino acids from water (1–5). These differentials in solvation energetics are remarkably large in magnitude (6,7), but to deform the lipid bilayer is also inherently costly (8–10). Therefore, the morphology of the protein-lipid interface ultimately reflects a trade-off between opposing energetic gains and penalties. The sensitivity of this balance, in turn, provides the cell with a means to regulate membrane protein function through mechanical inputs and variations in lipid content. Examples include diverse processes such as ion channel gating, protein topology, and oligomerization (11–13), even for proteins whose biological function is seemingly unrelated to membrane morphology.

Understanding the energetics of lipid bilayer deformations has therefore been the subject of decades of theoretical, computational, and experimental research. The first membrane shape Hamiltonian was developed by Helfrich, Canham, and Evans (8,9,14) in which the membrane is considered as a single 2D surface with intrinsic resistance to bending. Several extensions to the Helfrich continuum model improved its applicability to a broader range of biological phenomena. For example, phenomena like lipid orientation (15,16) and bilayer thickness variations (17,18) were shown to be increasingly important at molecular scales most relevant for membrane proteins (<100 Å), while other extensions addressed the effect of external forces like membrane stretch (19,20) on membrane shape. Each of these additions improved the agreement between theory and experiment or computation of thermal undulations in membranes, but whether they could be applied to deformations of significant magnitude, such as those created by integral membrane proteins, remained unexplored.

These protein-induced deformations were precisely the focus of Mouritsen and Bloom (21), Huang (10), and Wiggins and Phillips (22), among others. Mouritsen and Bloom focused primarily on hydrophobic thickness mismatch between embedded proteins and the membrane; their mattress model discretizes the membrane into a lattice of springs. Hydrophobic thickness mismatch is also a major component of the models proposed by Huang and Wiggins and Phillips, but these authors also address the membrane curvature component of localized thickness mismatch, to various resolutions: Huang returns to a continuum description of the membrane to describe gramicidin A dimerization, while Wiggins and Phillips’ bilayer-inclusion model for MscL gating uses a single “line tension” term that scales with the circumference of the channel to describe the energetic cost from the protein-lipid interface. Ultimately, this body of work provides parsimonious analytical models to explain the observed equilibria of protein systems; however, they also forgo a molecular, bottom-up derivation of the energetics of these larger-magnitude membrane shape distortions. (Of note, an orthogonal approach extracts the free energy of the membrane from the lateral pressure profiles, in systems with or without protein inclusions (23–25), but these studies do not address the effect of lateral tension specifically on deformed membranes).

The strengths of these continuum formulations of the membrane lie in their conceptual elegance and the distillation of a complex system into a few empirical parameters, such as the bending and area compressibility moduli, which can then be experimentally and computationally validated. For many cases of interest, however, the necessary descriptors are often not measurable or verifiable. One glaring example is heterogeneity in lipid composition, which is a fundamental aspect of biological systems. Heterogeneous lipid bilayers are challenging to model at the continuum-theory level because membrane shape and the spatial distribution of lipids are interdependent (26); no general analytical theory that we are aware of can anticipate with accuracy how different lipid species will distribute around a protein. Another complication is simply the variety of protein shapes and associated membrane perturbations. The assumptions of rotational symmetry or negligibility of area (10) or curvature and thickness terms (25) may become unjustifiable for more complex protein systems and lipid compositions. More fundamentally, whether composition or tension affect membrane energetics of large, thermally inaccessible curvature and thickness deformations remains to be systematically examined.

Computer simulations with molecular-scale resolution such as Molecular Dynamics (MD) have their own shortcomings, but as computing capacity and capability continue to increase, there is little doubt that simulations will remain the theoretical method of choice to examine the complex interplay between proteins and their surrounding membrane. The principal challenge that MD simulation studies face is the limited timescale of the processes that may be reliably examined, especially as the dimensions of the system under study reach the biological scale. A range of methodologies have been developed to overcome this limitation, however, collectively referred to as enhanced-sampling techniques (27,28). These approaches are designed to induce or increase the probability of rare events of interest. In the case of biological membranes, the Multi-Map method (29), which we use here, makes it possible to characterize the energetics of membrane deformations of any shape or scale, with any choice of lipid composition (30,31). Unlike continuum-mechanics models, the free-energy landscapes obtained through this approach reflect the complex structural dynamics and interatomic forces recorded in simulation, without requiring theoretical assumptions.

Here, we leverage both conventional simulations and enhanced-sampling simulations with the Multi-Map method to examine the effect of applied lateral stretch on free-energy landscapes of curvature and thickness deformations. We first recapitulate the result that tension has negligible effects on elementary mechanical properties of the membrane (17,18), then quantify the energetics of more extreme curvature and thickness deformations, allowing us to test the continuum theory at these higher energy states with and without tension. We further examine the effects of applied lateral stretch on lipid dynamics, asking whether it appreciably alters lipid mixing, diffusivity, and spatial localization. Finally, we assess how an altered lipid composition influences the morphological energetics of the membrane.

## Methods

The MD simulations in this study were computed with either GROMACS 2018.8 (32) or NAMD 2.14 (33), as noted. In all cases, simulations were based on the MARTINI force field, version 2.2 (34). In-house code was developed to make the MARTINI 2.2 parameter and topology files compatible with NAMD, available at https://github.com/1004parky/martini2_in_namd.

### Construction and equilibration of molecular systems

To examine how the spontaneous shape fluctuations of a membrane at rest change under stretch, we used lipid bilayers of 200 or 5,000 POPC molecules constructed for a previous study (29), whose lateral dimensions are 78 × 78 Å^2^ and 390 × 390 Å^2^, respectively. To examine how tension influences the energetics of induced deformations, we generated two additional bilayers, featuring different lipid compositions. These bilayers were constructed using the *insane* protocol (35) and consist of 400 lipids per leaflet (800 lipids total, 155 × 155 Å^2^), solvated with 9:1 water:antifreeze-water (34). The compositions of these bilayers are either 100% POPC or 50% POPC and 50% DLPC, distributed symmetrically between the two leaflets. The number of water particles for the pure POPC system is 22,944 and for the POPC/DLPC system is 23,753.

The two newly constructed 800-lipid bilayers were energy-minimized with GROMACS using the steepest-descent algorithm, for 500,000 steps. To begin to equilibrate these systems, we carried out three sequential MD simulations of 1 ns each, also in GROMACS, increasing the integration timestep from 2 to 10 to 20 fs; the temperature was set at 300 K through velocity-rescaling (36) and the pressure was set at 1 atm with the Berendsen semi-isotropic barostat (37). A cut-off value of 12 Å was used for both electrostatics and van der Waals interactions. After the third step, a fourth equilibration was carried out with NAMD, for 100 ns, also at 300 K and 1 atm, using a timestep of 25 fs and 12-Å cutoff for all nonbonded interactions. Unless otherwise noted, all the subsequent simulations used to evaluate the energetics and dynamics of the membrane in the microsecond timescale were calculated under the same conditions and energy functions. Representative simulation parameter files for the above steps (*.mdp for GROMACS, *.in for NAMD) are available on Zenodo (38).

### Simulation of applied lateral tension

In this work, we use “mechanical tension” (39,40) or “applied lateral stretch” (41) to refer to the outward stretching force applied to a lipid bilayer in the direction parallel to its midplane. (Others have studied the same force using slightly different terminology (9,11,17,20,22,25)). Importantly, this force is distinct from the persistent interfacial effect that exists at the lipid-water boundary, often referred to as “surface tension” or “interfacial tension” (39,40). To stretch the membrane laterally in simulation, we set the target pressures along the membrane plane, *P_xx_*^target^ and *P_yy_*^target^, to a value that differs from that along the membrane perpendicular, *P_zz_*^target^. The applied mechanical tension γ relates these three target pressures according to the following relationship:

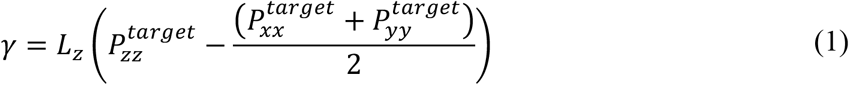

where *L_z_* is the length of the simulation box along the membrane perpendicular, and *P*_zz_ ^target^ = 1 atm. Note also that the target pressures *P_xx_*^target^ and *P_yy_*^target^ are equal when the simulation system is isotropic on the XY plane, as is the case in this study. To enforce these target pressures, we used the Langevin-piston method built into NAMD, using a 2-ps period and 1-ps decay. The 200-lipid and 5,000-lipid bilayers were simulated both with and without tension, with several values of γ ranging from 0 to 10 mN/m (1 mN/m = 1 dyn/cm); the 800-lipid bilayers were simulated at tension values of 0 and 10 mN/m.

### Definition of Multi-Map collective variables for different membrane perturbations

To induce different shape perturbations in the membrane and to calculate the associated free-energy profile in each case, we used the Multi-Map method (29) in combination with the umbrella-sampling technique (42). Specifically, we used two collective variables ξ_upper_ and ξ_lower_ to independently describe and define the morphology of each bilayer leaflet, through volumetric maps that represent their shape both at rest and at different magnitudes of a deformation of interest. For computational simplicity, only the lipid phosphate groups were considered in the definition of these collective variables; however, this choice did not artificially disrupt the integrity of the lipid bilayer (**Figure S4**). The variable used as reaction coordinate in the umbrella-sampling calculations, ξ, is the sum of ξ_upper_ and ξ_lower_.

To define an undeformed reference state of the membrane, we used volumetric maps that take the form of 1D Gaussian functions along the direction perpendicular to the membrane plane. Since both lipid composition and lateral tension influence the thickness of the bilayer at rest, the center of each Gaussian function was uniquely specified for each simulation system, based on analysis of the equilibration trajectory for that system (**Figure S1**). Because the phosphate groups naturally reside within the width of these Gaussian functions when the membrane is at rest, this design ensures that application of an umbrella potential on the Multi-Map variable does not deform the membrane, nor does it incur a free-energy cost, as is required for a valid reference state.

To induce a global bending deformation of membrane and to quantify its energetics, a set of volumetric maps were generated by convoluting the reference-state 1D Gaussian functions with a series of 1D sinusoidal functions defined as:

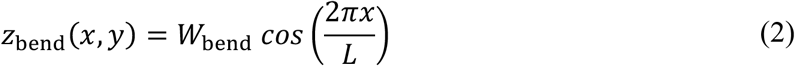

where *L* is the side-length of the membrane, and *W*_bend_ is equal to −10, −6, −2, +2, +6, or +10 Å (**Figure 1A**). As mentioned above, these 6 functions were used to define a Multi-Map variable for each leaflet, ξ_upper_ and ξ_lower_; their sum, ξ_bend_, was then used as the umbrella-sampling coordinate.

**Figure 1.**
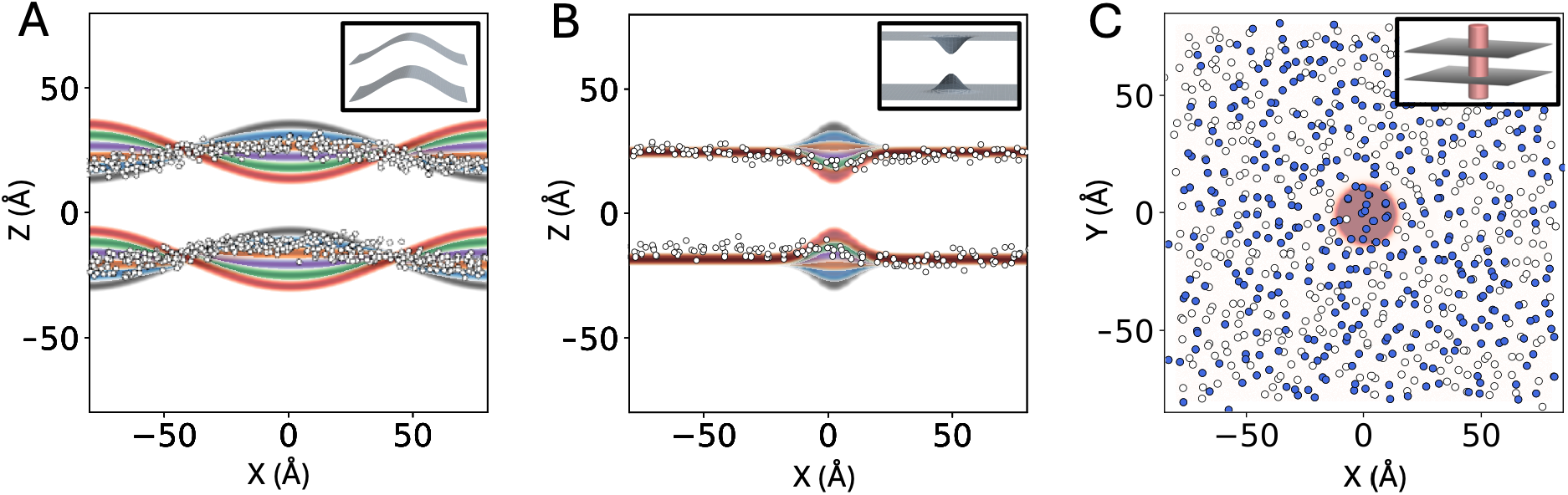
Volumetric maps used to induce and quantify perturbations in membrane shape and lipid distribution through Multi-Map simulations. (**A**) A cross-section of the set of maps used to study the energetics of a sinusoidal bending deformation in a POPC membrane. Overlaid is a cross-section of a snapshot of the phosphate layers, for ξ_bend_ ≈ 10, with phosphate groups represented as white circles. The inset shows a 3D contour of one set of maps, for each of the bilayer leaflets. (**B**) A cross-section of the set of maps used to study the energetics of a localized thickness defect. Overlaid is a cross-section of a snapshot of the phosphate layers, for ξ_thickness_ ≈ 0.7, represented as in (A). The inset shows a 3D contour of one set of maps. (**C**) A cross-section of the cylindrical map used to examine lipid dynamics. Overlaid is an XY-projection of a snapshot of the phosphate layers, for ξ_occupancy_ ≈ 15. Lipids designated as POPC_special_ are marked in blue. The inset shows a 3D contour of the target cylindrical volume, intersecting both leaflets of the bilayer.

To induce a localized thickness defect, the reference-state 1D Gaussian functions were convoluted with a series of 2D Gaussian curves of the form:

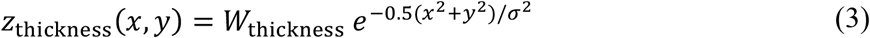

where *W*_thickness_ is equal to -10, -6, -2, +2, +6 or +10 Å, and σ determines the area affected by the thickness defect (**Figure 1B**). Specifically, we set σ = 8 Å such that the defect is about 40 Å in diameter. Analogously to the global deformation, these six maps were used to define a Multi-Map variable for each leaflet, ξ_upper_ and ξ_lower_, and these were combined into an overall reaction coordinate, ξ_thickness_. When defining ξ_upper_ and ξ_lower_, however, maps with opposing sign of

*W*_thickness_ were considered for the upper leaflet and lower leaflet; therefore, negative values of ξ_thickness_ correspond to membrane thickening (both leaflets bulging away from the midplane), while positive values reflect membrane thinning (both leaflets squeezing toward the midplane).

Finally, we used the Multi-Map method to induce the enrichment or depletion of a subset of the lipids in the membrane in a specific region, as a means to examine the impact of lateral stretch on lipid dynamics. Specifically, this subset of lipids, referred to as POPC_special_, are 50% of the POPC molecules in each leaflet, selected as random. The region targeted is a cylindrical volume of radius *R* of 10 Å, whose axis perpendicular to the membrane, spanning beyond both leaflets (**Figure 1C**). To induce the enrichment or depletion of POPC_special_ in this target region, we defined a Multi-Map variable ξ_occupancy_ that quantifies the total occupancy of the cylindrical volume by those lipids. For a given simulation snapshot, we defined the occupancy for each lipid *i* in the POPC_special_ subset as:

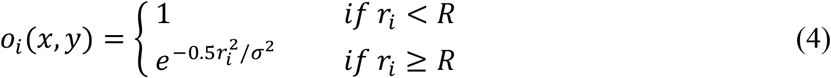

where *r_i_* denotes the distance from lipid *i* to the center of the cylinder, projected on the membrane plane. ξ_occupancy_ is therefore the sum of *o_i_* over all POPC_special_ lipids. Note that that occupancy function in Eq. 4 decays smoothly for *r_i_* ≥ *R*, specifically as a Gaussian of width σ = 2 Å. This design ensures that ξ_occupancy_ is continuous and differentiable. As with the other membrane perturbations described above, enhanced sampling simulations based on ξ_occupancy_ enabled us not only to control the population of POPC_special_ lipids in the target cylindrical volume, but also to quantify the associated free-energy gain or cost.

### Umbrella-sampling simulations using Multi-Map and interpretation of free-energy profiles

As mentioned, we used the umbrella-sampling method (42) to calculate the potential of mean force (PMF) along the reaction-coordinate defined by each of the Multi-Map variables ξ described above. To do so, we employed the Colvars module (43,44) interfaced with NAMD (33). In all cases, we initialized the umbrella-sampling windows with coordinates sequentially extracted from a 600-ns steered-MD trajectory in which a moving harmonic restraint was applied onto ξ, increasing gradually over 500 ns, with force constant of 5 kcal/mol. Once the initial coordinates were obtained, each window was simulated independently, with sampling times ranging from 1 to 7 µs per window; the first 300-ns fragment in each window was however considered equilibration time and excluded from the calculation of free energies. The weighted-histogram analysis method (WHAM) (45) was used to analyze the rest of the trajectories and derive the PMF as a function of the value of ξ in each case. For each of the PMF profiles discussed in this study we ascertained that the sampling times per window exceed the convergence time of the PMF (**Figure S2**).

The global bending deformation (**Figure 1A**) was induced by sampling values of ξ_bend_ from 0 to 20, in a series of 41 windows spaced by 0.5. The localized thickness deformation (**Figure 1B**) was induced by sampling values of ξ_thickness_ from -1.15 (thickening) to 1 (thinning), in a series of 44 windows spaced by 0.05. The enrichment or depletion of POPC_special_ was induced by sampling values of ξ_occupancy_ (**Figure 1C**) from 0 (depletion) to 16 (enrichment), in a series of 17 windows spaced by 1. The force constants of the static umbrella-potentials used in these calculations were 5 kcal/mol for ξ_bend_ and ξ_occupancy_ and 500 kcal/mol for ξ_thickness_. Note that because the volumetric maps that underlie ξ_bend_, ξ_thickness_, and ξ_occupancy_ are defined in absolute Cartesian space, all umbrella-sampling simulations also included a harmonic restraint applied to the center of mass of the bilayer, only along the Z-coordinate (membrane perpendicular), to ensure that the lipid bilayer remains in the frame of reference where the maps were constructed. This additional restraint has no effect on the results of the calculations, as the energy function used in our simulation systems is invariant to translations of the reference frame.

By definition, the collective variables ξ_bend_, ξ_thickness_, and ξ_occupancy_ are continuous and derivable functions of the atomic coordinates, as is required for any component of the energy function used in an MD simulation. However, while ξ_occupancy_ is an intuitive descriptor (it measures a number of molecules), ξ_bend_ and ξ_thickness_ are dimensionless quantities that might not be generally intuitive. Therefore, the free-energy profiles originally calculated using ξ_bend_ or ξ_thickness_ as reaction-coordinates are shown instead as functions of two physical descriptors that, although computed *a posteriori*, are linearly correlated with ξ_bend_ and ξ_thickness_ (**Figure S3**). These descriptors are the peak curvature of the membrane, *c*, in lieu of ξ_bend_, and the hydrophobic thickness of the membrane at the center of the defect, *t*, in lieu of ξ_thickness_.

To derive the peak curvature of the membrane for a given simulation snapshot, we first calculated the map of the total curvature *c*(*x*, *y*) that best fits the map of the membrane midplane, ℎ(*x*, *y*). Because the deformation depends only on the *x* coordinate (Eq. 2), the following expression can be used:

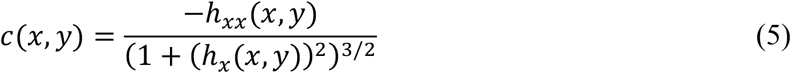

where the *x* suffix indicates a derivative along that component. We defined the mean value of *c* in each umbrella-sampling window as the average of the 1% largest values in the corresponding *c*(*x*, *y*) map. A linear regression between the mean values of ξ_bend_ and the mean values of *c* for each of the umbrella-sampling windows used in the calculation of PMF(ξ_bend_) produced an analytic relationship between ξ_bend_ and *c*, which we used to translate PMF(ξ_bend_) into PMF(*c*) (**Figure S3A**).

To quantify the hydrophobic thickness of the bilayer defect in a given simulation, we evaluated the mean distance between the outer and inner ester-group layers in an area within a radius of 0.04 × (box length) from the center of the defect (**Figure S5C**). We evaluated the bulk thickness *t*_b_ analogously, but in an area outside a radius of 0.3 × (box length) from the center of the defect, and within a radius of 0.425 × (box length) from that center (**Figure S5C**). To translate PMF(ξ_thickness_) into PMF(*t*), or PMF(*t* – *t*_b_), we again used the relationship between these variables deduced from a linear regression between the mean values of ξ_thickness_ and the mean values of *t* for each of the umbrella-sampling windows used in the original calculation of PMF(ξ_thickness_) (**Figure S3B, S3C**).

### Comparison of free-energy profiles to continuum-mechanics models

Free-energy profiles, computed numerically for each deformation, were compared to the corresponding theoretical predictions. For the sinusoidal bending deformation, the predicted free-energy cost as a function of the local curvature *c_local_* = *c*_1_ + *c*_2_ is 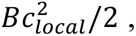 where *B* = *k_c_k_t_*/(*k_t_* + *k_c_q*^2^), *k_c_* and *k_t_* are the bending and tilt moduli for the bilayer, respectively, and *q* is the wavenumber of the sinusoid. The corresponding energy density expressed in terms of the peak curvature of the sinusoid, *c*, is *Bc*^2^*L*^2^/4, where *L* is the length of the bilayer.

For the Gaussian-shaped thickness deformations, we decomposed the model’s energy density into one term that accounts for the change in local thickness, and one term that accounts for the local curvature of the two leaflets, as done in previous work (10,46–49). The local thickness term was estimated by converting the thickness strain into the corresponding area strain and using the area compressibility modulus: *K_A_* ∑(*t*_Δ_*ν*_Δ_)^2^/2, where *t*_Δ_ = (*t* − *t_b_*)/*t_b_* and *ν*_Δ_ = (1 − *ν*)/*ν*. Here, *t* is the local thickness, *t_b_* is the bulk thickness, and *ν* is the Poisson ratio. Note that *ν* = 0.5 is typically used for bilayers, assuming volume incompressibility (50); however, since this assumption does not appear to hold for our simulated systems (**Figure 2A-B**), we estimated *ν* = 0.425 from the relative changes in area and thickness upon application of lateral tension. For the local curvature term, we summed the bending energy density of each leaflet calculated as 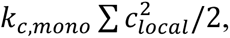 where *k_c_*_,*mono*_ = *k_c_*/2 estimates the bending modulus of one leaflet from the bilayer’s bending modulus. In this case, *c_local_* was determined for each leaflet separately, as the local curvature of the bilayer was 0 due to symmetry across the membrane midplane. Each term of the energy density was integrated over the projected two-dimensional surface of the membrane (**Figure 5A**).

**Figure 2.**
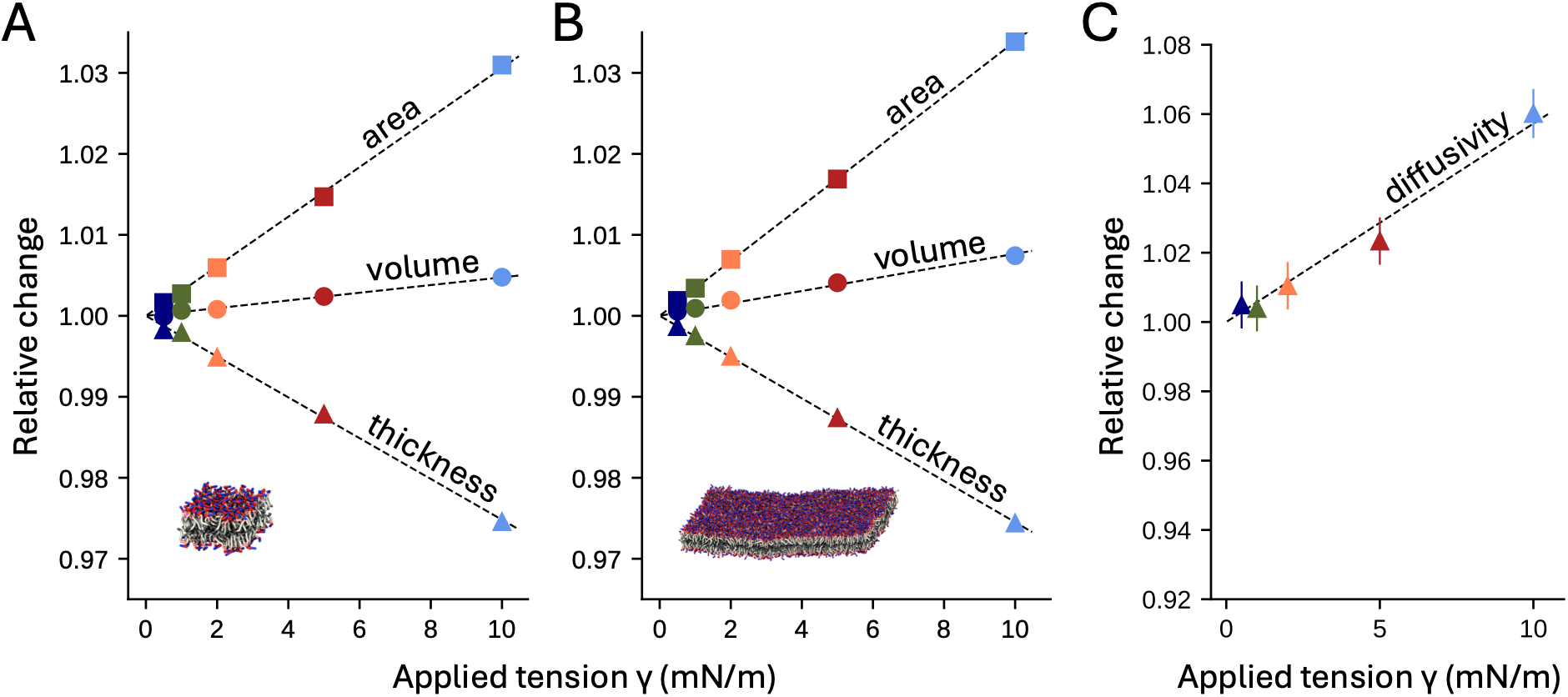
Effects of applied lateral stretch on membranes at rest. (**A**, **B**) Changes in area per lipid (squares), hydrophobic thickness (triangles), and volume (circles) caused by application of lateral tension of increasing magnitude to bilayers of 200 (A) or 5,000 (B) lipids. All values are in reference to the zero-tension condition. For each membrane and each tension condition, the values plotted derive from analysis of about 100 µs of sampling; the estimated statistical errors are smaller than the size of the symbols. (**C**) Changes in diffusivity *D,* for the 5,000-lipid bilayer, upon application of lateral tension of increasing magnitude, relative to the zero-tension condition. For each condition, *D* was deduced by calculating the linear slope of the relationship MSD = 2*D*τ, where MSD is the mean-squared displacement of all lipids in the membrane after an elapsed time τ in the 0 to 1 µs range; sampling errors were estimated from the spread of MSD values over lipid molecules.

All elastic moduli (*k_c_*, *k_t_*, and *K_A_*) were obtained from about 100 µs of sampling from 200- and 5,000-lipid membranes simulated at rest; their values are discussed in the Results.

### Evaluation of membrane structure and dynamics

The MOSAICS software suite (51) was used to analyze several descriptors of membrane structure and dynamics as observed in our simulations. Specifically, the *zcoord*, *midplane*, and *delta_plot* programs were used to calculate the membrane midplane and thicknesses based on the coordinates of the lipid ester groups (GL1/2). Lipid enrichment was evaluated with the *2d_enrichment* program. Each of these programs produced a 2D projection of the quantity analyzed across the plane of the membrane, for a designated simulation time-window. Lipid diffusion and kinetics were evaluated with the *lipid_mixing* program, which reports the fraction of lipids *j* that reside in the vicinity of lipid *i* (e.g. the first solation shell) for longer than a certain elapsed time (10 ns in our work). Fluctuation spectra were calculated with MDAnalysis (52) and fitted against theoretical predictions as summarized in the Results section; statistical uncertainties for fitted parameters were bootstrapped using 50,000 samples around the computed points (53).

## Results

### Effect of applied lateral stretch on membranes at rest

We started by studying the effect of lateral tension (γ, Eq. 1) on our systems by examining several key descriptors of membrane structure and dynamics, namely area per lipid molecule, volume, thickness, and lipid diffusivity. The basis for this analysis is a series of conventional MD simulations of both small and large POPC bilayers (200 and 5,000 lipids, respectively) with and without applied tension, approximately 100 µs in length for each value of γ. All the descriptors examined vary linearly with the magnitude of the applied tension (**Figure 2A**, **2B**). Intuitively, lateral stretch causes the area per lipid to increase and the membrane thickness to decrease. In particular, the area per lipid increases at a relative rate of 0.31% per mN/m for the 200-lipid bilayer, and 0.34% per mN/m for the 5,000-lipid bilayer. The inverse of these rates of change provides an estimate of the area compressibility modulus *K_A_* for our POPC bilayers, ranging from 322 ± 38 mN/m for the 200-lipid bilayer to 294 ± 11 mN/m for the 5,000-lipid bilayer, both of which are consistent with previously reported values for POPC membranes, obtained both experimentally (54,55) and computationally (55,56).

The changes in area per lipid outpace the changes in thickness, and so we observe an increase in membrane volume as the applied tension increases, and therefore a reduced lipid density. The increase in tension also results in an increase in lipid diffusivity, at a rate of ∼0.6% per mN/m (**Figure 2C**). This change appears to reflect the abovementioned increase in the area per lipid: lipid diffusion is a Brownian process that requires adjacent lipid molecules to exchange their relative positions on the membrane plane. As the area per lipid increases, the cumulative displacements that result from these exchanges also become larger, and hence the value of the diffusion constant *D* is also increased (**Figure 2C**).

Next, we examined two important mechanical properties of the membrane, namely the bending modulus, *k_c_*, and the tilt modulus, *k_t_*. Both descriptors can be derived from analysis of a membrane at rest, or more specifically, from analysis of the spontaneous fluctuations of the membrane midplane. Following the Monge parameterization (57), the membrane midplane is defined as the function *h*(*x*, *y*), where *h* denotes a vertical displacement relative to a flat plane. The ground state for a symmetric bilayer with no preferred curvature therefore corresponds to *h*(*x*, *y*) = 0. Thermal fluctuations, however, will naturally cause deviations from this idealized state, even for a membrane at rest (**Figure 3A**). In this regime, the bending and tilt moduli contribute to define the likelihood of a given shape fluctuation, through the following expression of the energy of the membrane (8,9,14–16,19):

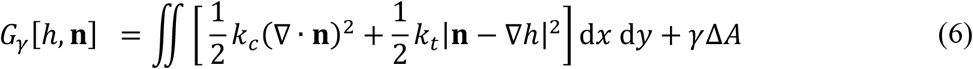

where **n**(*x*, *y*) is a position-dependent vector describing the local lipid orientation across the XY plane, ∇·**n** is the divergence of that vector, and ∇*h* is the gradient of the *h*(*x*, *y*) function. Note that the *γ*Δ*A* term in Eq. 6 is only relevant when the membrane is under stretch (19,57). Here, Δ*A = A*_ground_ *– A* quantifies the decrease in the area of the membrane (projected onto the XY plane) when the shape of the midplane fluctuates away from the perfectly flat ground state (**Figure 3A**). Note also that Δ*A* is positive in our definition; under outward stretch, i.e. when γ > 0, Eq. 6 predicts that fluctuations that reduce the projected area incur an energy penalty.

**Figure 3.**
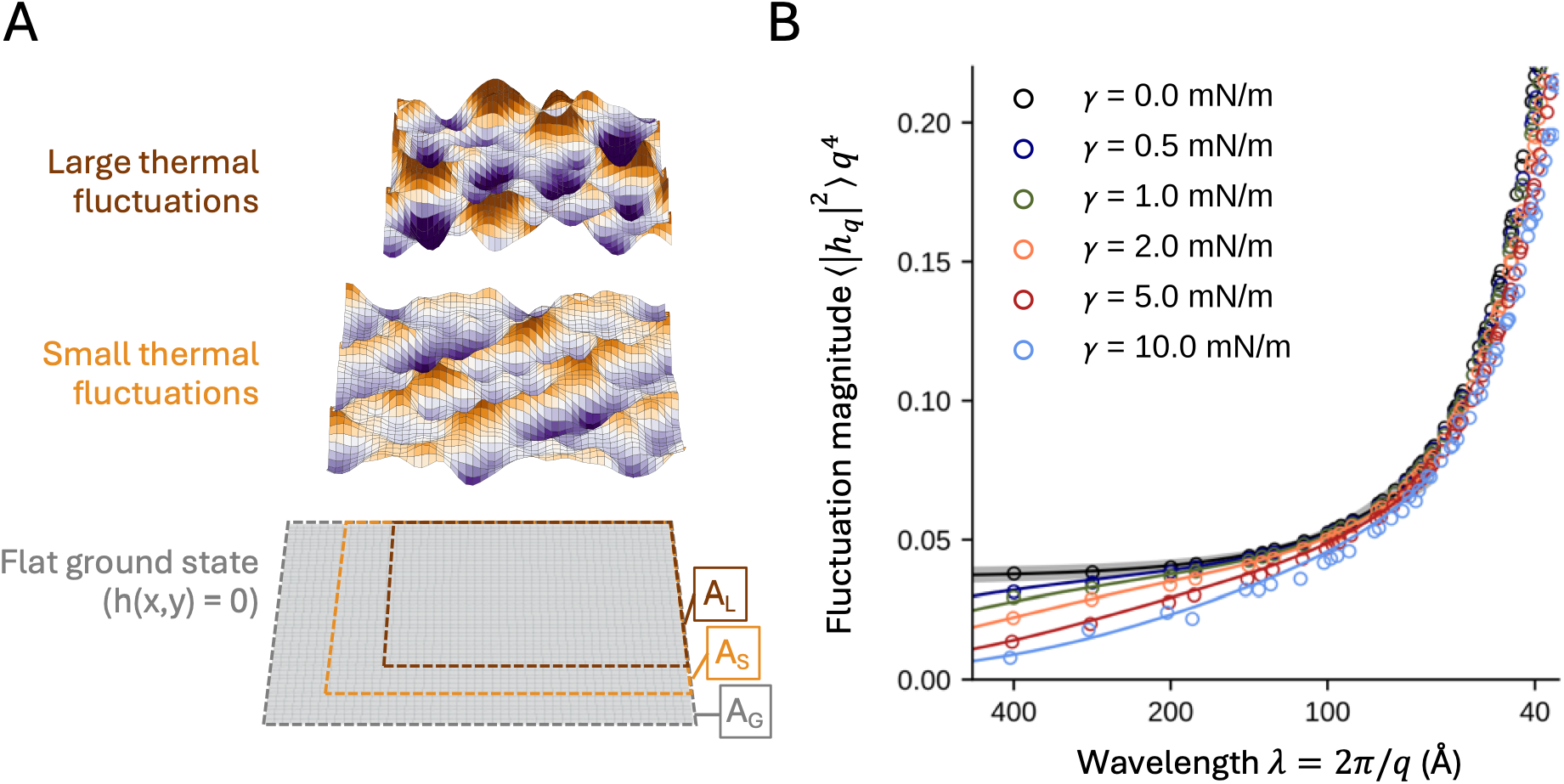
Impact of applied lateral tension on the spontaneous fluctuations of a bilayer at rest. (**A**) Illustrations of a membrane midplane in three different states, randomly generated in Python: a perfectly flat ground state, and two others with small and large thermal fluctuations, with color intensity (purple or orange) indicating larger amplitudes. Note how spontaneous thermal fluctuations naturally decrease the projected area of the membrane, from A_G_ (grey) to A_S_ (orange) to A_L_ (brown), respectively. Fluctuation modes such as these coexist at room temperature in the absence of stretch. However, upon application of lateral stretch, changes in projected area incur an energetic cost, given by γ(A_G_ – A_S_) or γ(A_G_ – A_L_) (Eq. 6). Fluctuations of larger magnitude and wavelength are thus increasingly unlikely to occur spontaneously under applied tension. (**B**) Fluctuation spectra for a 5,000-lipid bilayer, calculated under different lateral tensions, derived from conventional MD simulations of approximately 100 µs for each condition. The magnitudes of the observed fluctuation modes are plotted as circles. The black line corresponds to the best fit of Eq. 7 against the simulation data with γ = 0, which results in *k*_c_ = 16.3 ± 0.7 kcal/mol and *k*_t_ = 0.15 ± 0.02 kcal/mol/Å^2^. The gray area around the black solid line denotes the 95% confidence interval. Each of the other solid lines is Eq. 7 with the corresponding value of γ and the values of *k*_c_ and *k*_t_ obtained for γ = 0, with no additional fitting.

At a temperature *T*, the energy function in Eq. 6 predicts a spectrum of fluctuations of *h* whose magnitude is given by:

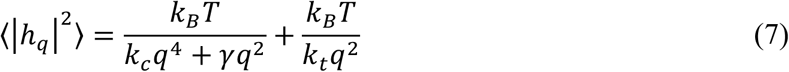

where *q* = 2π/*λ*, and *λ* is the wavelength of the fluctuation, *k_c_* is the bending modulus, and *k_t_* is the tilt modulus. The first term of Eq. 7 reflects the fluctuation spectrum predicted by the Helfrich theory under applied lateral tension (17,18,20,39,40), while the second term accounts for the fluctuations of the lipid orientation vector (15–17,20). Previous analyses of fluctuation spectra of MARTINI DPPC lipid simulations showed that both *k_c_* and *k_t_* changed by less than 10 % (close to their estimated error) under lateral tensions up to 20 mN/m (17). Thus, we examined |*h***_q_**|^2^ in our MD trajectories of the 5,000-lipid POPC bilayer and derived the values of *k_c_* and *k_t_* that result in the best fit between the simulation data and Eq. 7.

The spectrum for the unstretched bilayer (γ = 0) yielded values for both moduli with very good statistical precision, namely *k_c_* = 16.3 ± 0.7 kcal/mol, *k_t_* = 0.15 ± 0.02 kcal/mol/Å^2^ (**Figure 3B**), and in good agreement with previous studies of POPC bilayers under comparable conditions (58,59). Further, we found that no significant changes in either parameter were required to fit Eq. 7 onto the simulation data for any of the other tension conditions, up to γ = 10 mN/m (**Figure 3B**).

Because the lateral tension term in Eq. 6 depends on the projected area, it follows that the larger wavelength undulations are more heavily penalized by tension: 10 mN/m of applied tension suppresses the magnitudes of 400, 100, and 50 Å-wavelength fluctuations by 76%, 13%, and 2%, respectively. Or, expressed in terms of a free energy difference, 1.43 k_B_T, 0.14 k_B_T, and 0.02 k_B_T. Thus, from this analysis of thermal fluctuations alone, it is not evident how, or in what way, mechanical tension strongly influences the conformational energetics of membrane proteins that reside in this smaller wavelength, < 100 Å regime. A compelling hypothesis, of course, is that these proteins enhance their tension-sensitivity at this range by inducing membrane deformations beyond what thermal fluctuations permit, but it remains to be shown whether this γΔA term can be extended to these smaller-wavelength, larger-magnitude deformations associated with proteins.

### Effect of applied lateral stretch on global bending of the membrane

The first deformation that we examine beyond its thermally accessible magnitude is a sinusoidal deformation (**Figure 1A**). Using the Multi-Map simulation method, we quantified the associated free-energy change of deforming a 155-Å POPC lipid bilayer for amplitudes ranging from 0 to 10 Å (**Figure 4A**). Consistent with previous computations with Multi-Map (5,29,31) we observe that the energetic cost of bending the membrane increases with the magnitude of the induced curvature (**Figure 4B**). Remarkably, the Helfrich-based Hamiltonian (Eq. 6 with γ = 0, using the bending modulus *k_c_* and tilt modulus *k_t_* obtained from our simulations at rest) shows good agreement with our empirically determined potential of mean force (PMF), up to ∼ 20 k_B_T. However, as the magnitude of the deformation increases further, the free-energy profiles deviate significantly from that prediction (**Figure 4B**), possibly because the Helfrich-based models are quadratic approximations around the energy minimum. Our observations underscore why simulation-based free-energy calculations are invaluable in quantifying strongly deformed states.

**Figure 4.**
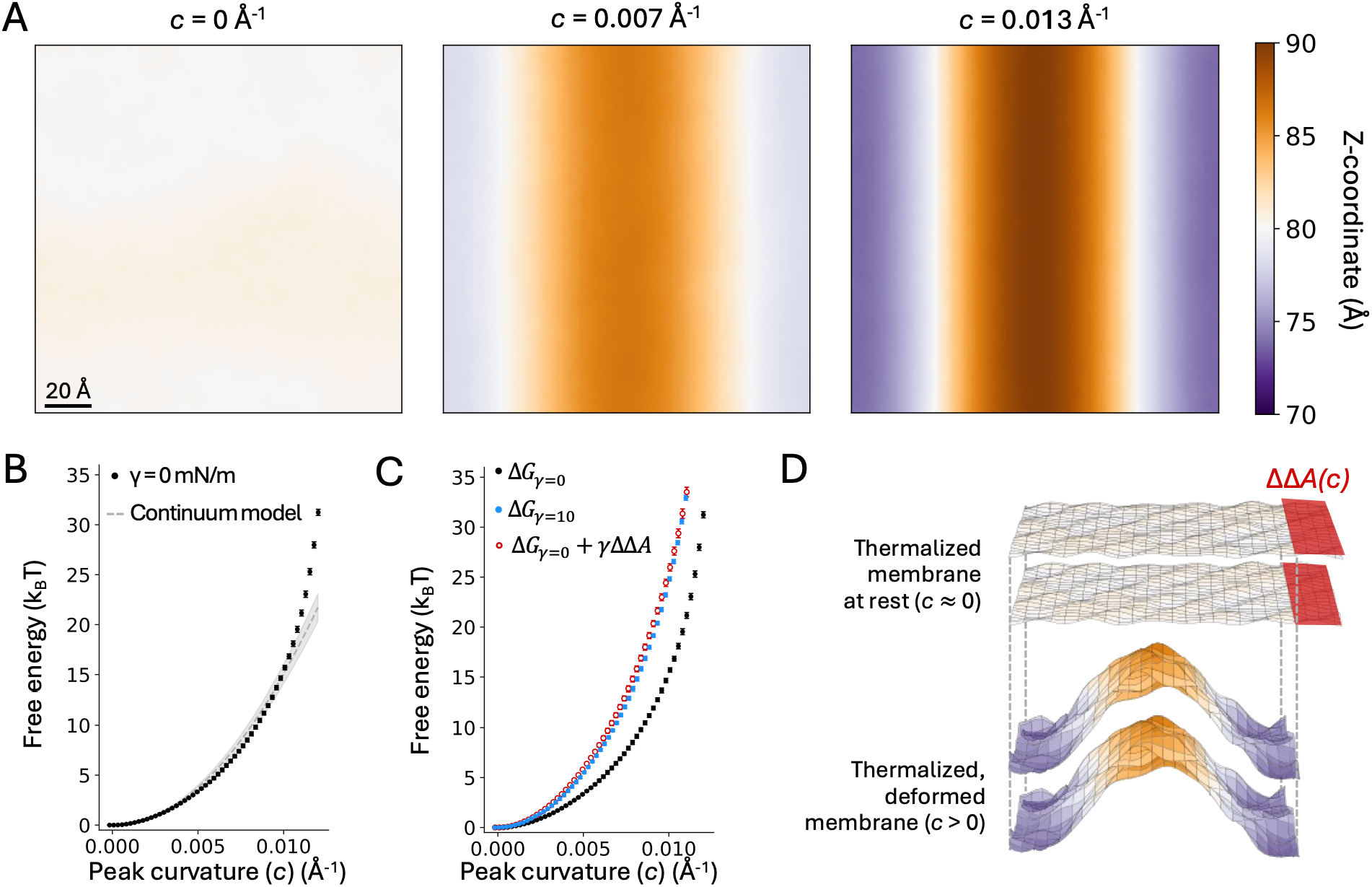
Effect of lateral tension on the energetics of membrane bending. (**A**) Two-dimensional maps of the membrane midplane for three different magnitudes of bending. (**B**) Potential of mean force (PMF) for a sinusoidal deformation of increasing curvature, under no tension. The free energy is plotted as function of the peak curvature of the deformation, *c*. The PMF curves were calculated with the Multi-Map method in combination with umbrella sampling, using an 800-POPC lipid bilayer. Each umbrella-sampling window was 1 µs. Fragments of 200 ns were used to estimate average values of the PMF (circles) and standard deviations, σ (bars); the error bars in the plot (σ) range from 0.002 to 0.19 k_B_T. The dashed line is the free energy predicted using continuum theory and the bending and tilt moduli derived from our simulations of membranes at rest. (**C**) Same data as in (B) but also showing the PMF of the same bilayer under applied tension (γ =10 mN/m, blue) and a computed curve (red) obtained by adding γΔΔA(*c*) to the PMF obtained with no applied tension (black). The errors in this additional profile, ranging from 0.004 k_B_T to 0.44 k_B_T, are the sum of the errors in the PMF calculated for γ = 0 and the errors in ΔΔA calculated with Eq. 8. (**D**) Schematic illustrating the changes in projected area of the membrane as a result of the sinusoidal deformation, with color intensity (purple or orange) indicating larger displacements relative to the monolayer midplane. ΔΔ*A*(*c*) is the difference in projected area between two thermalized states of the membrane, namely a reference, undeformed state with *c* ≈ 0 and a deformed state with *c* > 0. Note that the projected area of both states is smaller than that of a perfectly flat, idealized ground state.

Under the application of lateral tension at γ = 10 mN/m, we find that membrane bending becomes increasingly costly with respect to the peak curvature (see Methods) (**Figure 4C**). Further, this difference in energy between stretched and unstretched membranes can be entirely explained by a γΔΔ*A* term that reflects the change in the XY-projected area caused by the membrane deformation (**Figure 4D**). (We use *ΔΔA* rather than *ΔA* simply to acknowledge that both the stretched and unstretched membranes are thermalized at their respective tensions before being deformed.) For our sinusoidal deformation (Eq. 2), *ΔΔA* = 0.25*c*^2^*q*^−2^*L*^2^, where *c* is the peak curvature, *q* = 2*π*/*L* and *L* is the side-length of the membrane. The simplified form for the area change is:

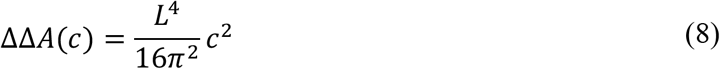

To derive *ΔΔA* from Eq. 8, we calculated the mean values of the peak curvature (*c*) and unit-cell length (*L*) in each of the umbrella-sampling simulations used to calculate the PMF profiles discussed above. Then, we added a term equal to γΔΔ*A*, with γ = 10 mN/m, to the PMF derived for the membrane without tension. As shown in **Figure 4C**, the resulting PMF is nearly identical to that directly calculated for the membrane under stretch. In addition, Eq. 8 reproduces the difference between the two PMFs well past the point where the PMF curves cease to be quadratic in *c*. This finding implies that for this type of membrane deformation, the increased stiffness of the membrane under lateral stretch is explained almost entirely by the degree of compression of its projected area. In other words, although the absolute free-energy of a bending deformation at ∼100- Å length scales may differ from Helfrich-based continuum theory at higher curvatures, the effect of lateral tension on bending energetics is completely consistent with a continuum treatment of the membrane.

### Effect of applied lateral stretch on localized thinning and thickening of the membrane

We next examined the effects of lateral tension on the energetics of a localized change in membrane thickness. To do so, we used the Multi-Map method to create and characterize Gaussian-shaped deformations of increasing magnitude, defined to be symmetric relative to the bilayer midplane (**Figure 1B**). For context, the characteristic Fourier wavelength (λ, **Figure 3B**) of this deformation is ∼50 Å, and we obtain PMFs for thicknesses ranging from 6 Å thinner and 6 Å thicker.

The time-averaged properties of the membrane, calculated from the z-coordinate and curvature maps of the two leaflets (**Figure S5**) as well as their thickness (**Figure 5A**), suggest that while the applied deformation alters both the local curvature and local thickness of the two leaflets, the larger deviations are in thickness; therefore, we plotted the PMF (**Figure 5B**) with respect to the thickness computed at the height of the deformation (**Figure S5C**). The PMF is not perfectly symmetrical, reflecting the fact that membrane thinning and thickening are not structurally equivalent processes. The comparison between our directly derived PMF and the continuum model (see Methods) is shown in **Figure 5B**. We show two predictions, one assuming volume incompressibility (yellow), and one where we estimate the Poisson ratio (grey) from the thickness and area compressions (**Figure 2**); both curves are reasonably similar to our PMF, although the inferred Poisson ratio certainly improves the agreement at higher magnitude deformations.

**Figure 5.**
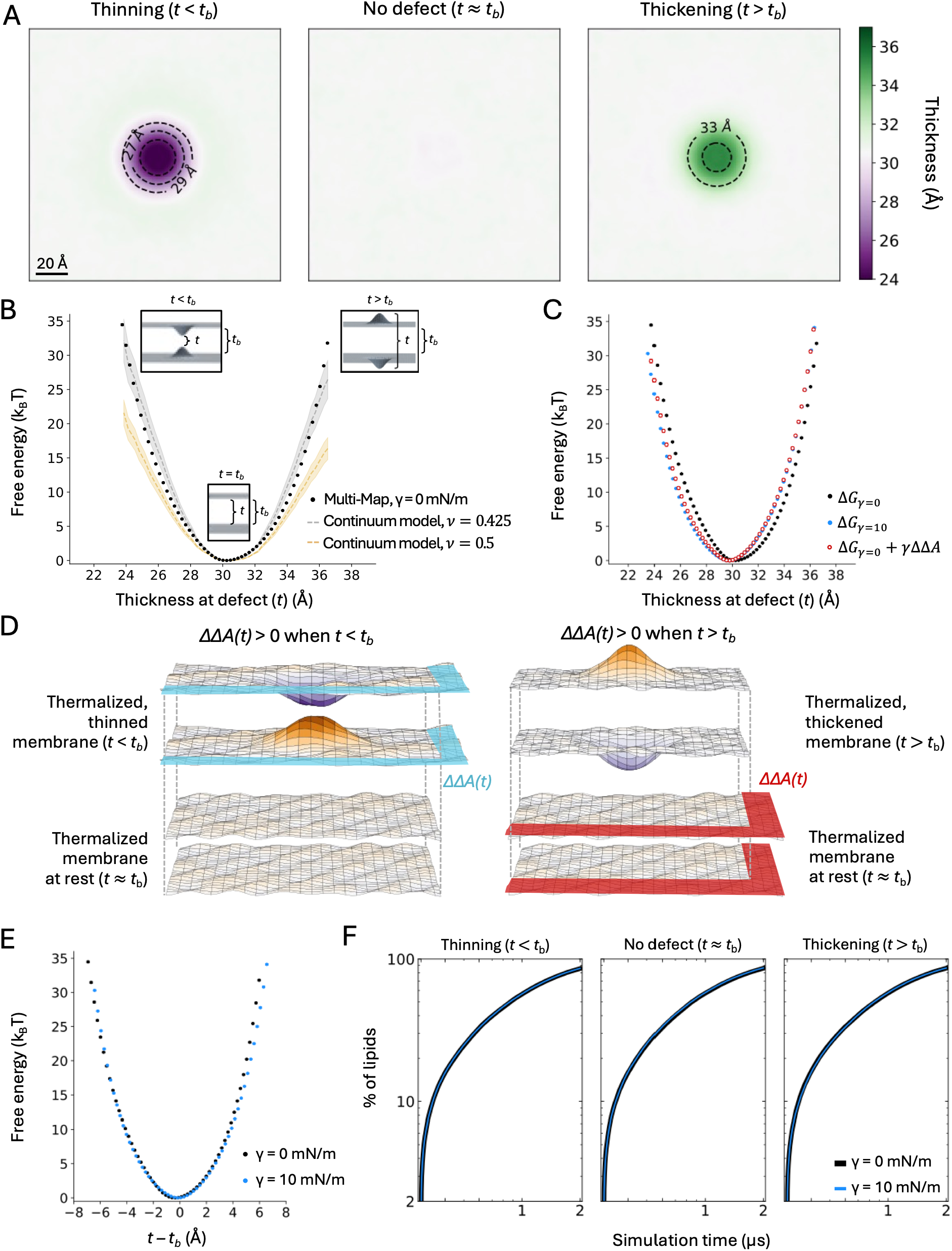
Effect of lateral stretch on the energetics of thickness perturbations. (**A**) Two-dimensional maps of the membrane thickness at three different deformation magnitudes. (**B**) Potential of mean force (PMF) for a localized membrane thickness defect, under no tension. The free energy is plotted as a function of the thickness at the center of the induced defect, *t* (black). The PMF curves were calculated with the Multi-Map method in combination with umbrella sampling, for an 800-POPC lipid bilayer. Each umbrella-sampling window was 2 µs. Fragments of 400 ns were used to estimate average values of the PMF (circles) and standard deviations, σ (bars); the error bars in the plot (σ) range from 0 to 0.13 k_B_T. The dashed lines are the predicted free energy from continuum theory, using the bending and compressibility moduli derived from our simulations of membranes at rest, and shown for two values of the Poisson ratio *ν*. (**C**) Same data as in (B) but also showing the PMF computed under applied tension (γ =10 mN/m, blue), and the corresponding predicted curve (red) obtained by adding γΔΔ*A*(*t*) to the PMF(*t*) profile calculated with no applied tension (black). The errors for this additional profile, ranging from 0.19 k_B_T to 0.26 k_B_T, are the sum of the errors in the PMF for γ = 0 and the errors in ΔΔ*A* calculated with Eq. 9. (**D**) Schematic illustrating the changes in the projected area of the membrane as a result of localized thickening or thinning, with color intensity (purple or orange) indicating larger amplitudes relative to the monolayer midplane. ΔΔ*A*(*t*) is the difference in projected area between two thermalized states of the membrane, namely a reference, undeformed state with *t* ≈ *t*_b_ and a deformed state with *t* > *t*_b_ or *t* < *t*_b_. Note that ΔΔ*A*(*t*) > 0 for *t* > *t*_b_, and ΔΔ*A*(*t*) < 0 for *t* < *t*_b_. (**E**) PMF curves with and without tension, as in (C), except the PMF data is plotted against *t* – *t*_b_, where *t*_b_ is the bulk thickness of membrane. (**F**) Evaluation of the impact of the thickness defect on the lipid dynamics. The plots quantify the degree of lipid mixing, which we define as the percentage of lipids *j* that reside for at least 10 ns in a 12 Å shell around each lipid *i*, for each leaflet. Full mixing is reached when this quantity is 100% i.e. all the possible lipid pairs *ij* have been observed to be first-neighbors for at least 10 ns. Note this condition is nearly achieved after 2 µs of simulation. Data is shown for three magnitudes of the thickness defect, namely *t* > *t*_b_, *t* ≈ *t*_b_, or *t* < *t*_b_, with or without applied tension γ = 10 mN/m. This comparison shows neither the thickness perturbation of the application of tension influences the lipid exchange dynamics.

Compared to the unstretched PMF, the PMF at γ = 10 mN/m is shifted to the left (**Figure 5C**). The structural reason for this shift is that lateral tension decreases the bulk membrane thickness *t*_b_ by ∼0.7 Å. Based on our findings for the global bending deformations and for the thermal fluctuations at rest, we reasoned that this differential effect might again stem from changes in membrane area induced by the thickness perturbation, and how those changes are penalized or favored under stretch. Here, we deduced the relative change in projected area caused by this perturbation, *ΔΔA*, from an evaluation of *t*, the thickness at the center of the defect. More specifically, we used:

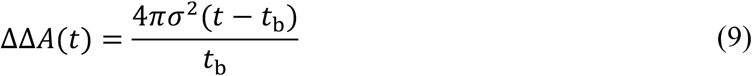

where *σ* = 8 Å is the half-width of the two-dimensional Gaussian function used to create the thickness defect, 2πσ^2^ is the integral of that function, and the additional factor of 2 accounts for the fact both bilayer leaflets are simultaneously perturbed. As for the bending deformation, we evaluated the mean value of *t* for each of the umbrella-sampling simulations used to generate the thickness defect without applied tension, and then calculated the additional free-energy term γ *ΔΔA*, with γ = 10 mN/m. Note that in this case *ΔΔA* is positive for the thickened defect (*t* > *t*_b_) but is negative for the thinned defect (*t* < *t*_b_) (**Figure 5D**). With γ > 0, the free-energy term γ*ΔΔA* therefore penalizes the former but favors the latter.

As shown in **Figure 5C**, when that γ*ΔΔA* term is added to the PMF calculated for the tensionless condition, the resulting curve is in excellent agreement with the PMF derived from MD simulations under γ = 10 mN/m. Thus, although lateral tension clearly affects the free-energy landscape of thickness deformations, it does so primarily through this γ*ΔΔA* term while preserving many of the other properties of the membrane. Specifically, the similarity in the shape of the free-energy curves calculated with and without tension when they are aligned by their respective minima (**Figure 5E**) suggests that for modest (< 5 Å) *relative* changes to the membrane thickness, mechanical tension does not significantly alter the resistance of the membrane to that change. In other words, as observed from analyses of thermal fluctuations, the bending and compressibility moduli appear to be largely independent of the range of lateral tension studied here.

Finally, it is worth clarifying that neither lateral tension nor the membrane thickness defects generated in this analysis have a discernable impact on the lipid mixing (**Figure 5F**). Thus, the modest tension-dependent changes in diffusivity (**Figure 2C**) do not appreciably impede the turnover of lipids at the membrane deformation.

### Does applied lateral stretch influence the dynamics of lipids in the membrane?

The results presented above demonstrate that lateral stretch influences the average structure of the membrane, the magnitude of its shape fluctuations at rest, and the energetics of morphological changes that might be induced extrinsically. Our results also show that neither mechanical tension, membrane thickening, nor membrane thinning appreciably alter the dynamics of lipid mixing. The implication is that proteins that thicken or thin the membrane can do so without affecting lipid turnover at those sites, and further that direct binding of lipid molecules would be a poor sensor for applied lateral tension.

To test this hypothesis using a protein-free model lipid bilayer, we used the Multi-Map method to induce the explicit enrichment or depletion of a specific subset of lipid molecules in a certain volume within the membrane, with and without applied tension, then evaluated in each condition the rate at which those lipids enter (and exit) the target volume. We also quantified the free-energy cost of this process of enrichment or depletion, and compared the result with what would be mathematically expected for a homogeneous membrane at rest. For simplicity, we used a cylinder of 10-Å radius as the target volume, which approximately corresponds to 8 lipids per leaflet (both with and without tension). The subset of lipids that were subject to enrichment or depletion comprise 50% of the membrane. These “special” lipid molecules, referred to as POPC_special_, were selected at random and are, of course, identical to the other lipid molecules. Therefore, any enrichment or depletion of POPC_special_ in the target volume leaves the structure and energy of the membrane unchanged, and the corresponding gains or losses in free energy would consist entirely of entropy.

As shown in **Figure 6A**, the Multi-Map simulations successfully sampled all possible occupancy states of the target volume, ranging from one in which no POPC_special_ lipids are found in the target volume (in which case only “normal” POPC lipids occupy the volume) to another in which the only lipids residing in the volume are POPC_special_ (see also **Figure S6A**). Logically, not all these states are equally probable; both with and without lateral stretch, the PMF curves reflecting the free-energy cost of this enrichment or depletion show that the most probable states are those in which 50% of the lipids in the target volume are “special” lipids, while “normal” lipids fill the rest. This is the expected result for a condition that preserves the natural dynamics of the membrane, since as mentioned, POPC_special_ comprises 50% of the lipids in the membrane. Indeed, not only the minima but the totality of the PMF curves derived from the Multi-Map simulations reproduce what would be mathematically predicted for a volume roughly equivalent to 8 lipids per leaflet, given that the membrane comprises 800 POPC lipids, 50% of which are designated as POPC_special_ (**Figure 6A**). That is:

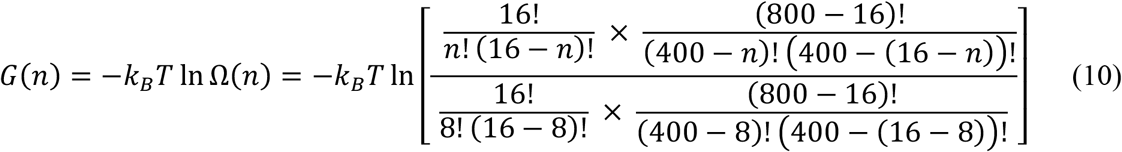

where *n* is the number of POPC_special_ lipids that reside in the target volume.

**Figure 6.**
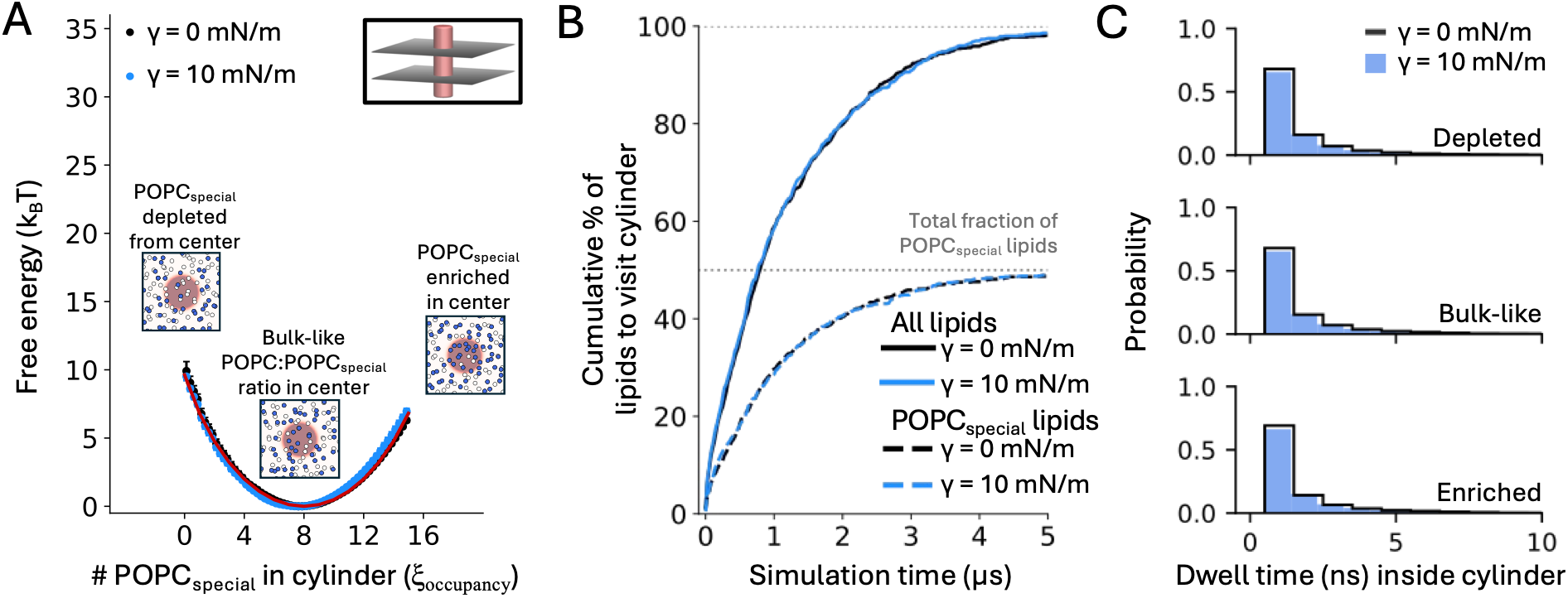
Applied lateral tension does not drive lipid dynamics. (**A**) Potential of mean force (PMF) for enriching or depleting lipids designated as POPC_special_ in a cylindrical volume of 10-Å radius at the center of the membrane, under no stretch (black) or under applied lateral tension of 10 mN/m (blue). The free energy is plotted as a function of the number of POPC_special_ lipids in the target area, ξ_occupancy_. The PMF data were calculated with the Multi-Map method in combination with umbrella sampling, for an 800-POPC lipid bilayer. Inset figures capture a snapshot of the simulation system in the umbrella-sampling windows calculated for ξ_occupancy_ values of 0, 7, and 15. Each umbrella-sampling window was 5 µs. Fragments of 1 µs were used to estimate average values of the PMF (circles) and standard deviations, σ (bars); the error bars in the plot (σ) range from 0.01 k_B_T to 0.69 k_B_T. The red line shows the PMF curve that is mathematically predicted purely based on entropic factors, using Eq. 10. (**B**) Cumulative percentage of lipids in each leaflet of the membrane that enter the target cylindrical volume after a certain simulation time, with (blue) or without applied tension (black), counting all lipids (solid lines) or only POPC_special_ lipids (dotted lines). These data derive from the umbrella-sampling window for ξ_occupancy_ = 7; analogous results were obtained for other values of ξ_occupancy_ (Figure S6). (**C**) Distribution of observed dwell-times of POPC_special_ lipids inside the cylinder, in each of the umbrella-sampling windows depicted in (A). Data obtained with applied tension (blue, filled) and without (black, unfilled) are compared. Bin widths for the two tension conditions differ for clarity of visualization.

We next quantified whether mechanical tension influences the rate at which lipids of either type enter (and exit) the target volume. As demonstrated in **Figures 6B** and **6C**, we detected no discernible effect on lipid dynamics, even at 10 mN/m. That is, the rate at which lipids of either kind entered the target volume, thereby replacing other lipids therein, was unchanged when the membrane was laterally stretched (see also **Figure S6B**). In both conditions, we observe that after 5 µs of simulation virtually every single lipid molecule in the bilayer, whether designated as POPC_special_ or not, has entered the target volume at least once, and dwelled therein for some time before exiting. As expected for Brownian motion, these single-molecule dwell times vary widely, from tens of nanoseconds to microseconds; yet, these distributions are also largely unchanged under lateral stretch (**Figure 6C**). In conclusion, we detect no evidence of a through-space “pulling” force influencing the dynamics of lipids that specifically results from the application of stretch. As demonstrated in the previous sections, lateral tension does influence the structure of the membrane and its conformational energetics, but its only effect on lipid dynamics is a minor increase in overall diffusivity.

### Localized thinning and thickening defects in a mixed membrane

The last system we examined was a mixed bilayer composed of a 1:1 ratio POPC and DLPC (di 12:0 PC) lipids in each leaflet. This is an interesting system to consider specifically for membrane thickness deformations because a 1:1 POPC:DLPC membrane is thinner, on average, than a pure POPC bilayer, owing to DLPC’s shorter alkyl chains. We reasoned, therefore, that shape perturbations in this mixed membrane, whose altered lipid composition leads to a smaller *t*_b_, might be energetically comparable to those in the homogeneous membrane under tension.

As mentioned above, application of lateral tension diminishes the bulk thickness of the membrane, *t*_b_; for POPC, in our simulation conditions, that shift is about 0.07 Å per mN/m of tension. In simulations of bilayers at rest, we find that the bulk-membrane thickness of the mixed bilayer is about 2.3 Å smaller than the pure POPC bilayer (**Figure 7A**). **Figure 7C** compares the PMF profiles for creating a localized thickness defect in pure POPC and in the 1:1 POPC:DLPC bilayer, without any tension applied. As we observed when contrasting pure POPC with and without applied tension, the PMF curve for the membrane with a smaller *t*_b_ shifts to the left. The magnitude of the shift caused by the addition of DLPC is larger than that caused by lateral stretch, consistent with the larger change in *t*_b_, the thickness at rest. As a result of this shift, the range of hydrophobic thicknesses that are realistically attainable are very different for the mixed bilayer than for pure POPC.

**Figure 7.**
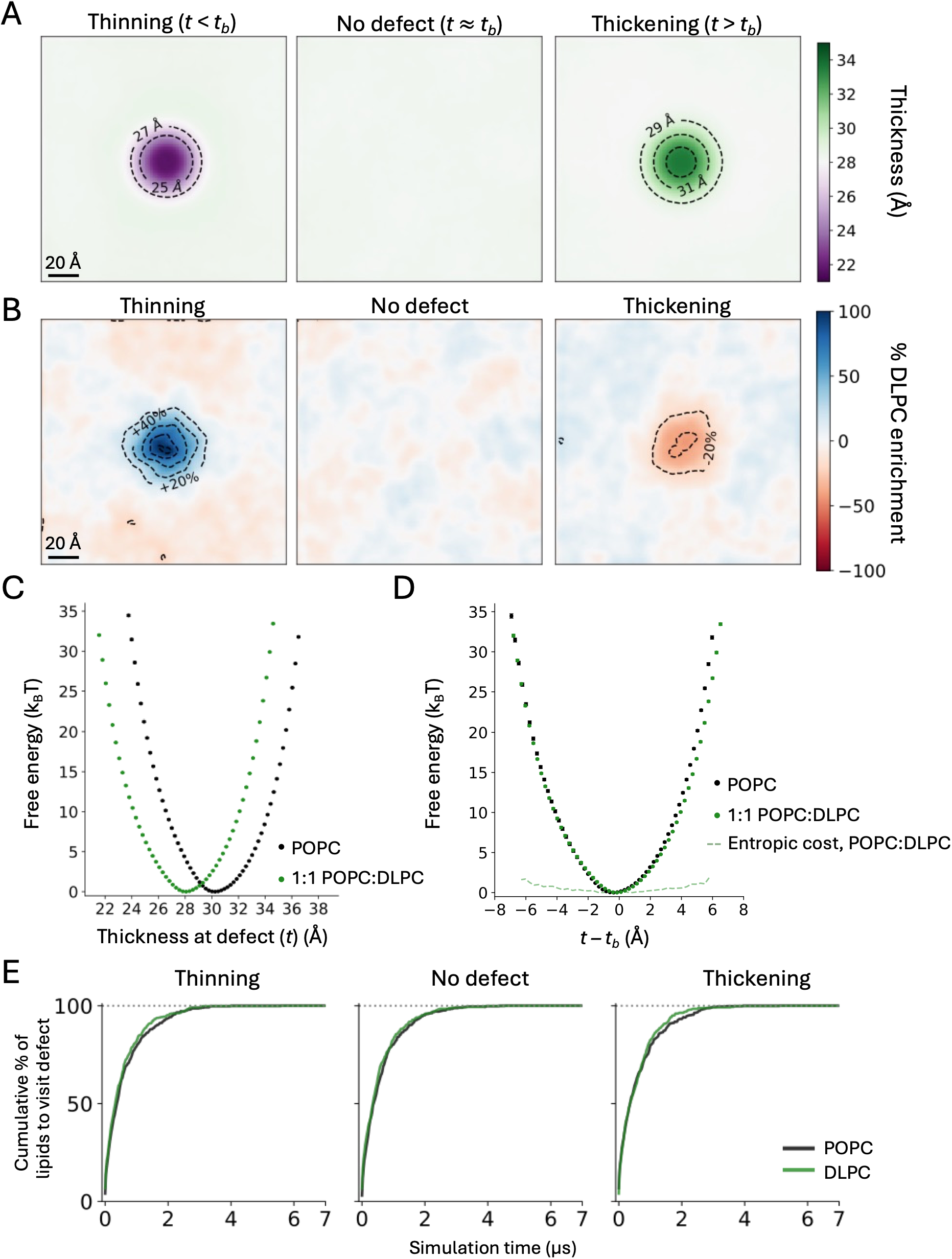
Influence of lipid composition on the energetics of thickness perturbations. (**A**) Two-dimensional map of the hydrophobic thickness of the mixed POPC:DLPC membrane for three different magnitudes of the induced central defect, namely *t* < *t*_b_ (left), *t* ≈ *t*_b_ (center) or *t* > *t*_b_ (right). Both POPC and DLPC lipid molecules contribute to define these maps. Dotted contour lines indicate 2-Å increments or decrements. The thickness value in the bulk, *t*_b_, is 28.2 Å. (**B**) Distribution of DLPC lipids in the three states represented in (A); an enrichment level of 0% corresponds to the bulk 1:1 POPC:DLPC ratio, positive enrichment values reflect accumulation of DLPC relative to that ratio, and negative values indicate depletion. Dotted contour lines indicate 20% increments or decrements. (**C**) Potential of mean force (PMF) for a localized membrane thickness defect. The free energy is plotted as a function of the hydrophobic thickness at the center of the induced defect, *t*, in POPC (black) or 1:1 POPC:DLPC (green) membranes. The PMF curves were calculated with the Multi-Map method in combination with umbrella sampling, for an 800-lipid bilayer with the indicated composition. Each umbrella-sampling window was 2 µs for the POPC membrane and 7 µs for the mixed membrane. Fragments of 400 ns (POPC) or 1.4 µs (1:1 POPC:DLPC) were used to estimate average values of the PMF (circles) and standard deviations, σ (bars); the error bars in the plot (σ) range from 0 to 0.13 k_B_T. (**D**) Same as in (C), except the PMF data are plotted against *t* – *t*_b_, where *t*_b_ is the bulk thickness of membrane. In addition, the entropic cost associated with the lipid enrichments in (B) is shown. This entropic cost constitutes just a fraction of the total PMF. (**E**) Cumulative percentage of POPC or DLPC lipids that enter the target cylindrical volume, in each leaflet of the membrane, after a certain simulation time, for each of the states represented in (A) and (B). Note that after 4 µs of simulation all lipids have visited the target volume, and that the turnover of DLPC lipids is slightly faster than POPC ones.

Interestingly, overlay of the PMF curves calculated for the mixed and pure membranes shows minimal differences at low deformation magnitudes, although there is a slight softening of the POPC:DLPC PMF beyond ∼5 Å thinning or ∼3 Å thickening (**Figure 7D**). Addition of DLPC therefore appears not to cause significant changes in the mechanical properties of the membrane, compared with pure POPC. It is worth noting that the spatial distribution of POPC and DLPC lipids is significantly changed by the introduction of the thickness perturbation. Specifically, in the thinned defect we observe a pronounced enrichment of the shorter-chain DLPC lipids, where the lipid content deviates from the bulk 1:1 ratio by as much as 80% (**Figure 7B**). Conversely, in the defect that thickens the membrane, DLPC lipids are depleted and POPC enriched (**Figure 7B**). These effects are entirely spontaneous, but not equivalent: DLPC enrichment in the thinned defect is stronger than its depletion in thickened defect. (The likely explanation is that the presence of POPC lipids in the thinned defect is strongly penalized because it causes steric clashes between opposing leaflets; by contrast, DLPC seems to be merely disfavored in the thickened defect, as it leads to a suboptimal degree of chain interdigitation across the midplane.) This partial lipid de-mixing naturally incurs an entropic energetic cost that is already incorporated into the Multi-Map computed PMF. We can, based on the average enrichment maps (**Figure 7B**), estimate the entropic contribution of the PMF (**Figure 7D**); at the most extreme deformations, it is about 2-2.5 k_B_T. Thus, although we observe a slight softening of the PMF at large deformations, the larger effect can be attributed to the shift in bulk thickness (**Figure 7C**) rather than a change in the mechanical properties of the mixed membrane.

Finally, it is worth emphasizing that the observed accumulation of POPC or DLPC in the induced thickness defect does not imply that individual lipid molecules are localized, or that the lipid dynamics is somehow arrested. In fact, we observed that after 4 µs of simulation time every single lipid molecule in the membrane had visited the thickness deformation at least once, for both the thinned and thickened defects (**Figure 7E**). As expected, DLPC diffuses slightly more quickly than POPC, generally (**Figure 7E**). However, the single-molecule dwell times inside the defect were nearly identical for the two lipid types (**Figure S7**), indicating that enrichment or depletion of a certain species at the defect reflects differential probabilities for entry into and exit from that locale, rather than a slow-down of the lipid dynamics. From a methodological standpoint, we would argue that these interdependent observations underscore one of the key advantages of the Multi-Map simulation framework. That is, through this approach it is possible to quantitatively evaluate how variations in lipid content influence the energetics of membrane morphology, while simultaneously assessing how changes in morphology influence the spatial distribution of different lipid types.

## Discussion

The morphological energetics of the lipid bilayer are undeniably influential for membrane protein structure and function, individually or in complexes. This influence has been however difficult to quantify, both experimentally and computationally, as protein structure or activity can hardly be evaluated without a membrane, or at least a medium that mimics it. Nonetheless, it is self-evident that the membrane is not a passive, infinitely adaptable medium; it features its own conformational preferences, both in terms of shape and lipid distribution. Integral membrane proteins must therefore contend with the lipid bilayer energetics; if a protein features alternate structural states that reshape the membrane or redistribute its lipid content differently, as has been demonstrated for a range of channels, transporters, and enzymes (1,5,11–13,41,60,61), the activity of that protein will be necessarily sensitive to extrinsic factors that alter the morphology or chemical composition of the membrane.

Mechanosensitive proteins are the most salient example of this interdependence. Well-studied ion channels like PIEZO, MscL, MscS, and their homologs belong to this class of proteins. Like other ion channels, these proteins naturally exist in an equilibrium between conducting (open) and non-conducting (closed) states. What differentiates these mechanosensitive channels is that this equilibrium shifts in response to applied lateral stretch, even in the absence of any other proteins or cellular components. This intrinsic mechanosensitivity has been described as the “force-from-lipids” model of mechanosensation (distinguishing it from “force-from-tether” or “force-from-filament” mechanosensitive proteins that rely on cytoskeletal connections) (62). The “force-from-lipids” model has been further subdivided into a “pulling” model, where tension is transduced to the channel via direct lipid-protein binding interactions (62–65); or an entropy-based or “delipidation” model, where tension promotes the outward movement of lipids in protein pockets and thereby enable structural rearrangements in the protein (4,62,66,67). An alternative framing of the “force-from-lipids” model explicitly places membrane energetics at the center of the gating mechanism: mechanosensitive channels sense lateral tension because the membrane deformations that stabilize their closed states become increasingly energetically costly with applied stretch (2,68).

That lateral tension affects the energy landscape of membrane deformations is well established; a large body of work has extended the Helfrich theory to describe the energetics of the membrane in terms of its curvature, lipid tilt, and mechanical tension (8–10,15–17,20,46,69). What our current work shows is that the continuum description of the effect of lateral tension on membrane shape deformations (namely, γΔΔA_proj_) remains appropriate for curvature and thickness deformations of significant magnitude. Interestingly, this γΔΔA_proj_ term accurately transforms the PMF obtained at zero tension to the PMF under tension, even though the Helfrich theory’s prediction is less accurate at large curvatures, potentially due to the quadratic approximations inherent to these models (**Figure 4A-B**). The γΔΔA_proj_ term is equally applicable for combined thickness and curvature deformations, whose zero-tension PMF is better described by continuum models (**Figure 5B-C**). Importantly, our conclusions do not rely on the assumptions specific to each theory: the free-energy profiles presented in this study stem solely from the intermolecular forces and preferred configurations observed during our MD simulations.

Precisely because our approach preserves the molecular behavior of the system, we were also able to directly examine the effect of lateral tension on the dynamics of lipid molecules. As noted above, two prominent models of mechanosensation rest upon lateral tension disfavoring lipid residence near the protein. A potential source of confusion may arise from the interdependence between the protein and membrane structure: protein-induced membrane deformations are indeed stabilized by interactions between the protein and the membrane, and mechanical tension may disfavor these interactions by penalizing the membrane deformations, leading to membrane flattening. Phenomenologically, this sequence of events can be rationalized under either the delipidation model or the membrane deformation model. However, our analysis of lipid dwell times and mixing in strongly deformed membranes (**Figure 5F**) or membranes where specific lipids are biased away or toward the center of the membrane (**Figure 6B-C**, **S5**) clearly indicates that applied tension does not exert an outward force on lipids. Our systems here are simulated without protein, but we also observed the same in systems with embedded proteins (2). Together, these studies clarify that lateral tension affects the conformational energetics of the membrane, rather than the lipid occupancy at certain sites. The membrane deformation model of mechanosensation is therefore consistent with an understanding of lateral tension that is rooted in physical principles. It also provides a rationale for why the force sensitivity of PIEZO, MscS, and MscL scales with the spatial extent and magnitude of their membrane perturbations (4,25,62).

The observations of constant lipid exchange also apply to a mixed membrane (**Figure 7E, S6D-E**), despite the stable enrichment of certain lipid species when a local defect is induced to the bilayer’s thickness. This is consistent with a preferential solvation mechanism (70), which has been previously demonstrated to explain the effect of membrane composition on protein dimerization (60). Together, these results suggest that altered lipid compositions which lead to altered enrichment profiles within membrane defects (or near proteins) can significantly shift the free-energy landscape without disrupting the dynamics and fluidity of the constituent lipids. Surprisingly, although we observe this free-energy shift for the mixed membrane as a function of absolute membrane thickness values (**Figure 7C**), the similarity in the shapes of the aligned PMF curves (**Figure 7D**) suggests that this lipid de-mixing does not significantly alter the elastic properties of the membrane, at least for the MARTINI 2.2 lipids used here. Our PMFs indicate a slight softening at > 5 Å membrane thickening or thinning, although within the estimated error in computed moduli. Future work on other lipid mixtures will certainly enrich our understanding of the role that preferential solvation plays in membrane deformation energetics and protein conformational landscapes.

In conclusion, we have demonstrated that lateral stretch alters the morphological energetics of lipid bilayers, supporting the notion that deformations in membrane curvature and thickness underlie the controlling effect of applied tension on the activity of mechanosensitive ion channels (2,41,68,71). We have also shown that tension has a marginal effect on lipid dynamics, indicating that it does not directly influence the lifetime of specific protein-lipid interactions that might occur. Clearly, much more ground remains to be covered to attain a comprehensive understanding of mechanosensation, and more broadly, of the growing number of biological processes known to entail membrane deformations of different magnitude and scale (12,13,71–73). We anticipate that molecular-simulation methods like Multi-Map (29) will continue to reveal important insights in areas of interest to membrane biophysics, such as for atomistic, asymmetric, or non-planar bilayers, provided these methods are designed to enable realistic modeling and quantitative evaluation of non-native bilayer states, both in terms of shape and lipid distribution.

## Data Availability

Source code for all MD simulations is publicly available as part of the Colvars library (https://colvars.github.io/) and the NAMD package (https://www.ks.uiuc.edu/Research/namd/). Initial and final simulation snapshots, as well as parameter input files, are freely available from Zenodo (38); to download full trajectory data (1.5 TB), please contact the authors. Python code developed specifically for this study can be freely downloaded from GitHub as noted above.

## Author Contributions

Conceptualization: YCP, GF, JDFG; methodology: YCP, GF; investigation: YCP, GF; analysis: YCP, GF; writing: YCP, GF, JDFG; supervision: GF, JDFG; funding acquisition: GF, JDFG.

## Acknowledgements

This study was funded by the Divisions of Intramural Research of the National Heart, Lung and Blood Institute (YCP, GF and JDFG), and of the National Institute of Neurological Disorders and Stroke (ZIA NS003143, GF). The authors are most thankful to Lucy R. Forrest for her support and sponsorship. All computational resources were provided by the National Institutes of Health High-Performance Computing Cluster.

