## Supplemental Figures for "On the effect of applied lateral tension on the deformation energetics of biological membranes and the lipid dynamics within"

July 23rd, 2026

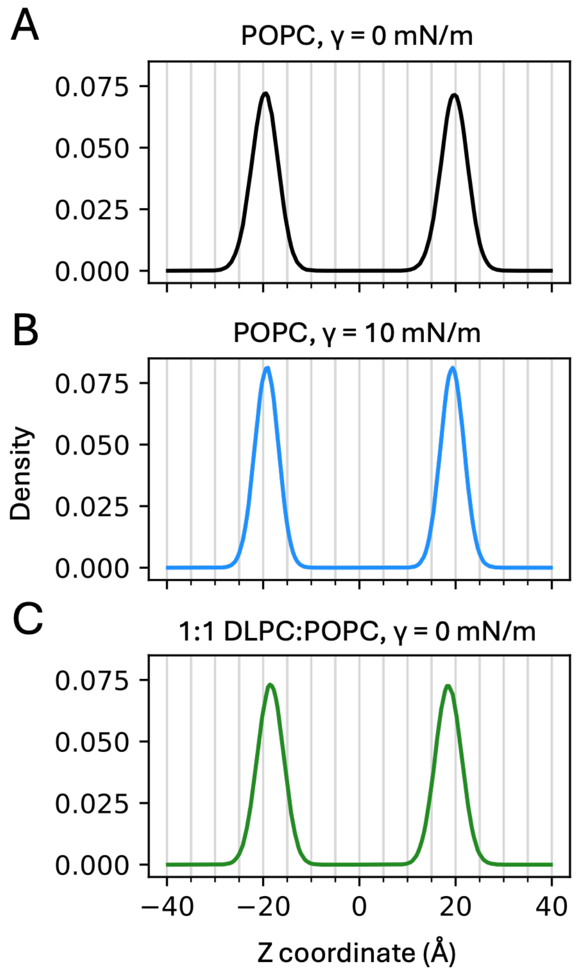


**Figure S1. Location of the phosphate layers in different simulation systems and conditions.** The figures show density profiles along the direction perpendicular to the membrane plane (Z) for the lipid phosphate groups. The profiles derive from analysis of 100-ns equilibration trajectories. The positions and width of the peaks observed in each case were used to define the reference, undeformed state of the membrane in each of the Multi-Map free-energy simulations (Methods). The density profiles fit 1D Gaussian functions with (**A**) $\mu = \pm 19.7$ Å and $\sigma = 2.8$ Å, (**B**) $\mu = \pm 19.3$ Å and $\sigma = 2.5$ Å, and (**C**) $\mu = \pm 18.5$ Å and $\sigma = 2.7$ Å.

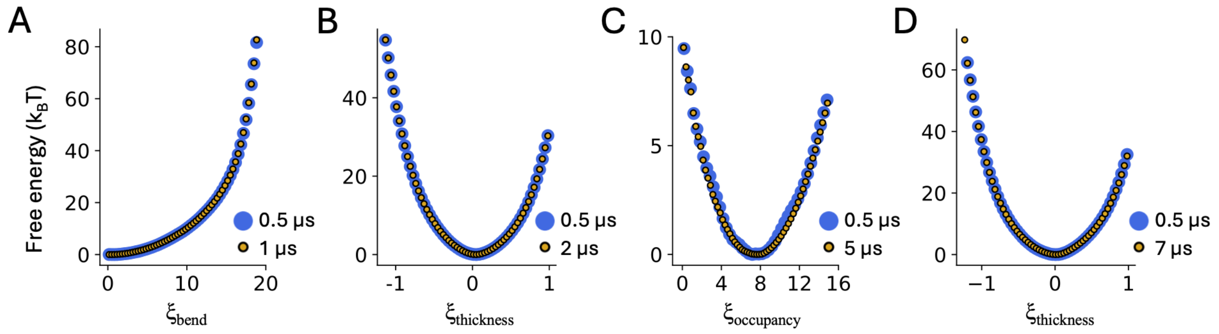


**Figure S2. Analysis of PMF convergence.** (**A**) Comparison of PMF profiles calculated for a sinusoidal deformation of a POPC membrane, under applied tension $\gamma = 10$ mN/m, after either 0.5 μs (blue) or 1 μs (yellow) of sampling time, demonstrating convergence. The latter is also shown in Figure 4. (**B**) Comparison of PMF profiles calculated for a localized thickness defect in a POPC membrane, with $\gamma = 10$ mN/m, after either 0.5 μs (blue) or 2 μs (yellow) of sampling time. The latter is also shown in Figures 5 and 7. (**C**) Comparison of PMF profiles calculated for the enrichment or depletion of $\text{POPC}_{\text{special}}$ at the membrane center, with $\gamma = 10$ mN/m, after either 0.5 μs (blue) or 5 μs (yellow) of sampling time. The latter is also shown in Figure 6. (**D**) Comparison of PMF profiles calculated for a localized thickness defect in a 1:1 POPC:DLPC membrane, after 0.5 μs (blue) or 7 μs (yellow) of sampling time. The latter is also shown in Figure 7.

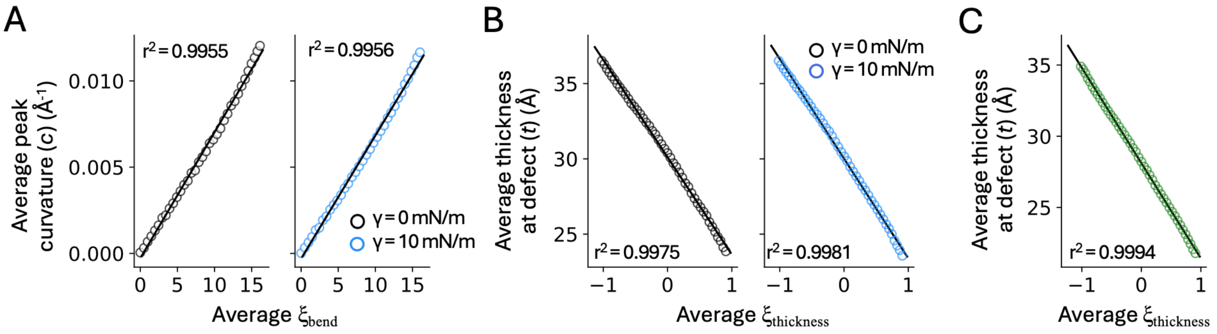


**Figure S3. Correlation between Multi-Map variables ξ and physical descriptors of membrane shape.** (**A**) The mean values of $\xi_{bend}$ recorded in each of the umbrella-sampling simulations used to induce a sinusoidal deformation (Figure 4) are plotted against the mean values of the peak curvature of the deformation, *c*. Data are shown for $\gamma = 0$ (left, black) and $\gamma = 10$ mN/m (right, blue). Linear regressions of these data (solid lines) result in $c = 0.00073\xi_{bend} - 0.00029$ for $\gamma = 0$ and $c = 0.00071\xi_{bend} - 0.00030$ for $\gamma =$ 10 mN/m. These relationships were used to translate PMF($\xi_{bend}$) into PMF(*c*) for each tension condition. (**B**) Same as (A), for the thickness deformation examined in Figure 5, i.e. mean values of $\xi_{thickness}$ are plotted against mean values of the hydrophobic thickness at the center of the defect, *t*. Data are also shown for $\gamma =$ 0 (left, black) and $\gamma = 10$ mN/m (right, blue). for $\gamma = 0$ (left, black) and $\gamma = 10$ mN/m (right, blue). Linear regression of these data (solid lines) result in $t = -6.46\xi_{thickness} + 30.11$ for $\gamma = 0$ and $t = -6.63\xi_{thickness} + 30.02$ for $\gamma = 10$ mN/m. These relationships were used to translate PMF($\xi_{thickness}$) into PMF(*t*) for each tension condition. (**C**) Same as (B), for the 1:1 POPC:DLPC condition (Figure 7). In this case, the linear regression results in $t = -6.69\xi_{thickness} + 28.12$.

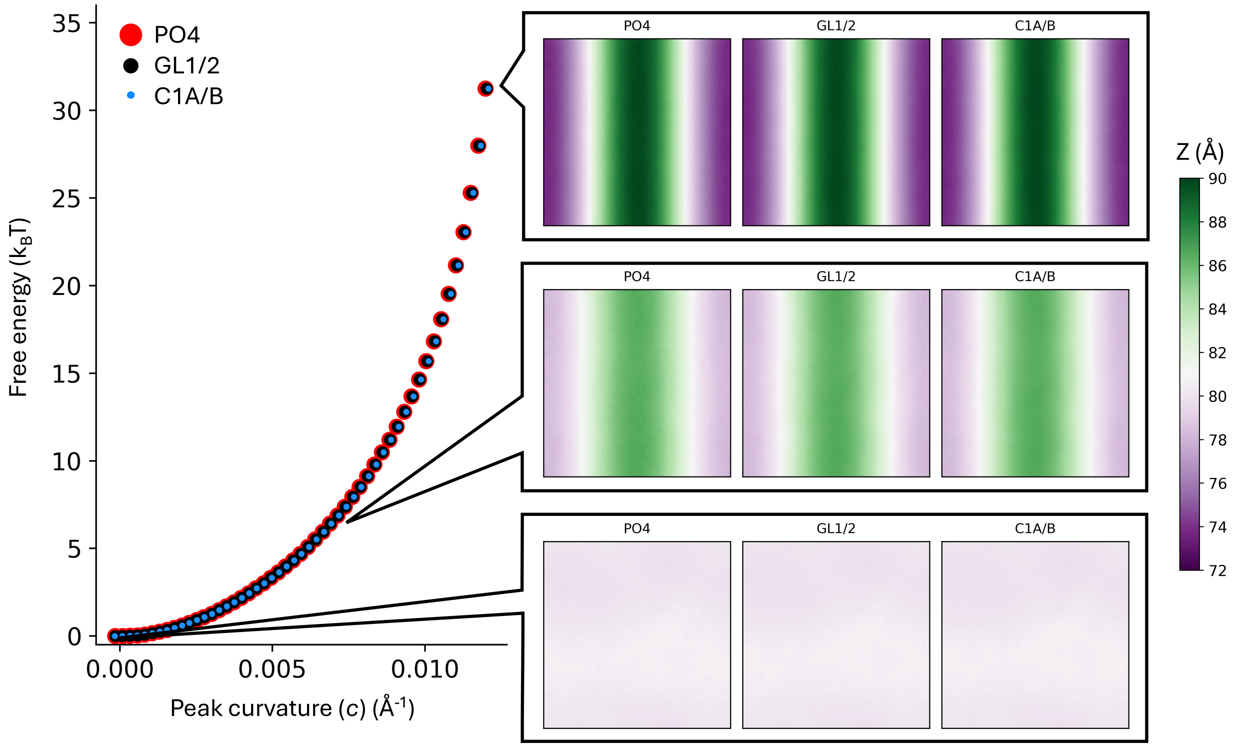


**Figure S4. Membrane structure is preserved across bead types analyzed.** The free energy of bending as a function of the peak curvature ($c$) obtained from midplane maps (right) generated from Z-coordinates of phosphate (PO4, red), glycerol (GL1/2, black), or first carbon tail (C1A/B, blue) beads. Insets show 2D midplane maps from three umbrella sampling windows for each bead type. Note that the GL1/2 PMF is the same as in Figure 4B, and all 2D maps and PMFs are virtually identical across bead types.

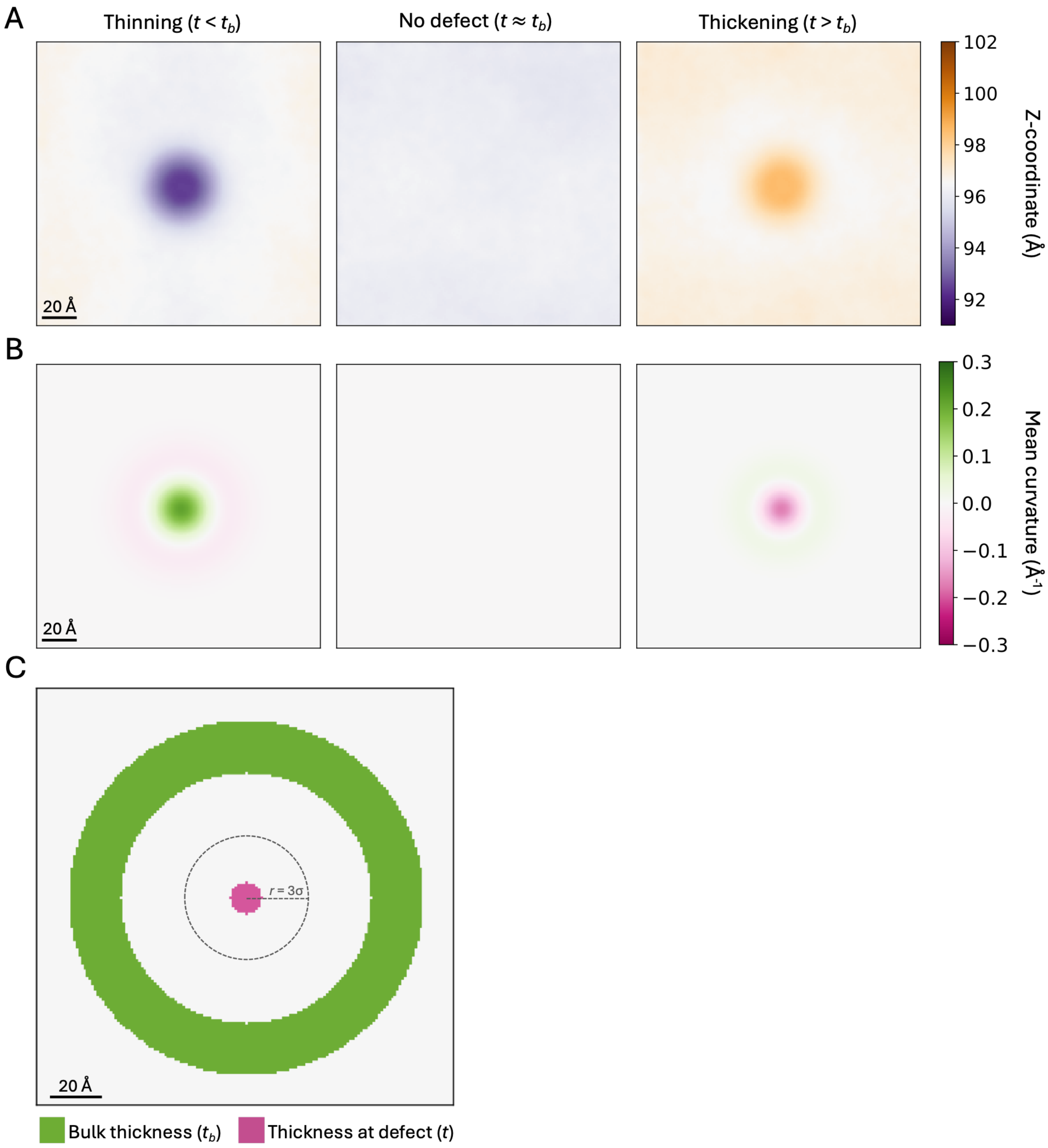


**Figure S5. Analysis of Gaussian deformation simulations**. (**A**) Two-dimensional maps of the z-coordinate of the upper leaflet for thinning (left), no defect (middle), or thickening (right). (**B**) Two-dimensional maps of the mean curvature $(c_1 + c_2)/2$ of the upper leaflet for the windows indicated in (A). (**C**) Masks used to evaluate the separation between the two ester layers either at the center of the thickness defect ($t$, pink) or in the bulk ($t_b$, green). Values within each region were averaged to obtain a single value of $t$ or $t_b$ for each system or simulation condition. Regions in white were not considered. In particular, the outermost shell was not considered to ensure that the edges of the simulation box, where lipids move between periodic boundaries, do not skew the analysis. The dotted lines indicate the extent of the deformation, where $\sigma = 8$ Å is the half-width of the Gaussian function used to create the thickness defect.

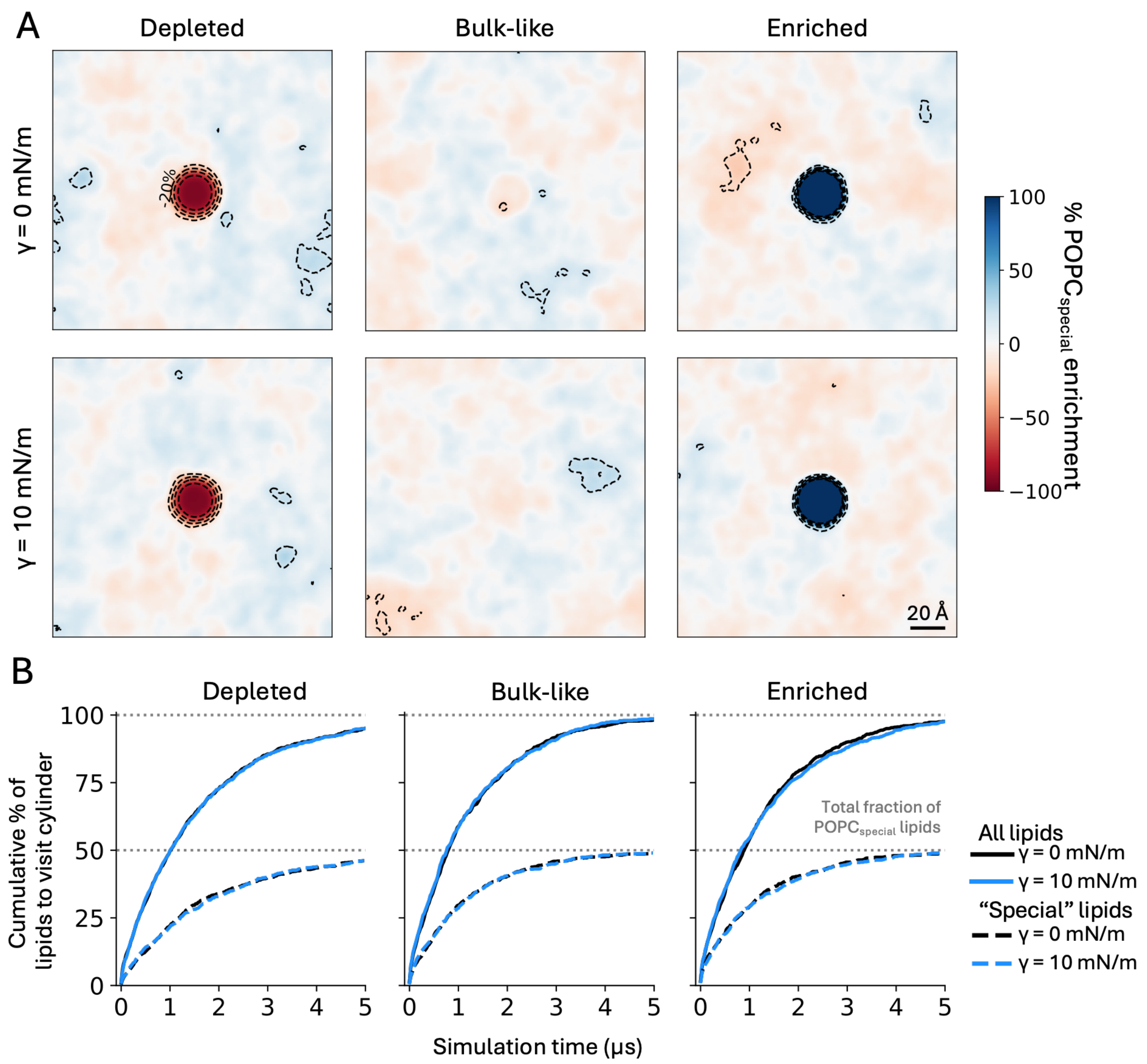


**Figure S6. Applied tension has no impact on lipid exchange dynamics.** (**A**) Distribution of $POPC_{special}$ lipids in the Multi-Map simulations associated with Figure 6, with for $\xi_{occupancy} = 0$ (fully depleted), $\xi_{occupancy} = 7$ (bulk-like), or $\xi_{occupancy} = 15$ (fully enriched) after 5 μs of simulation. An enrichment level of 0% corresponds to the bulk ratio of $POPC_{special}$ i.e. 50%. Dashed contour lines indicate enrichment increments of 20%. (**B**) Cumulative percentage of lipids in each leaflet of the membrane that enter the target cylindrical volume after a certain simulation time for each of the states represented in (A), with (blue) or without tension (black), counting all lipids (solid lines) or only $POPC_{special}$ lipids (dotted lines). Note that after 5 μs of simulation, nearly all lipids have visited the target volume, and that the application of tension has no appreciable effect on the lipid dynamics.

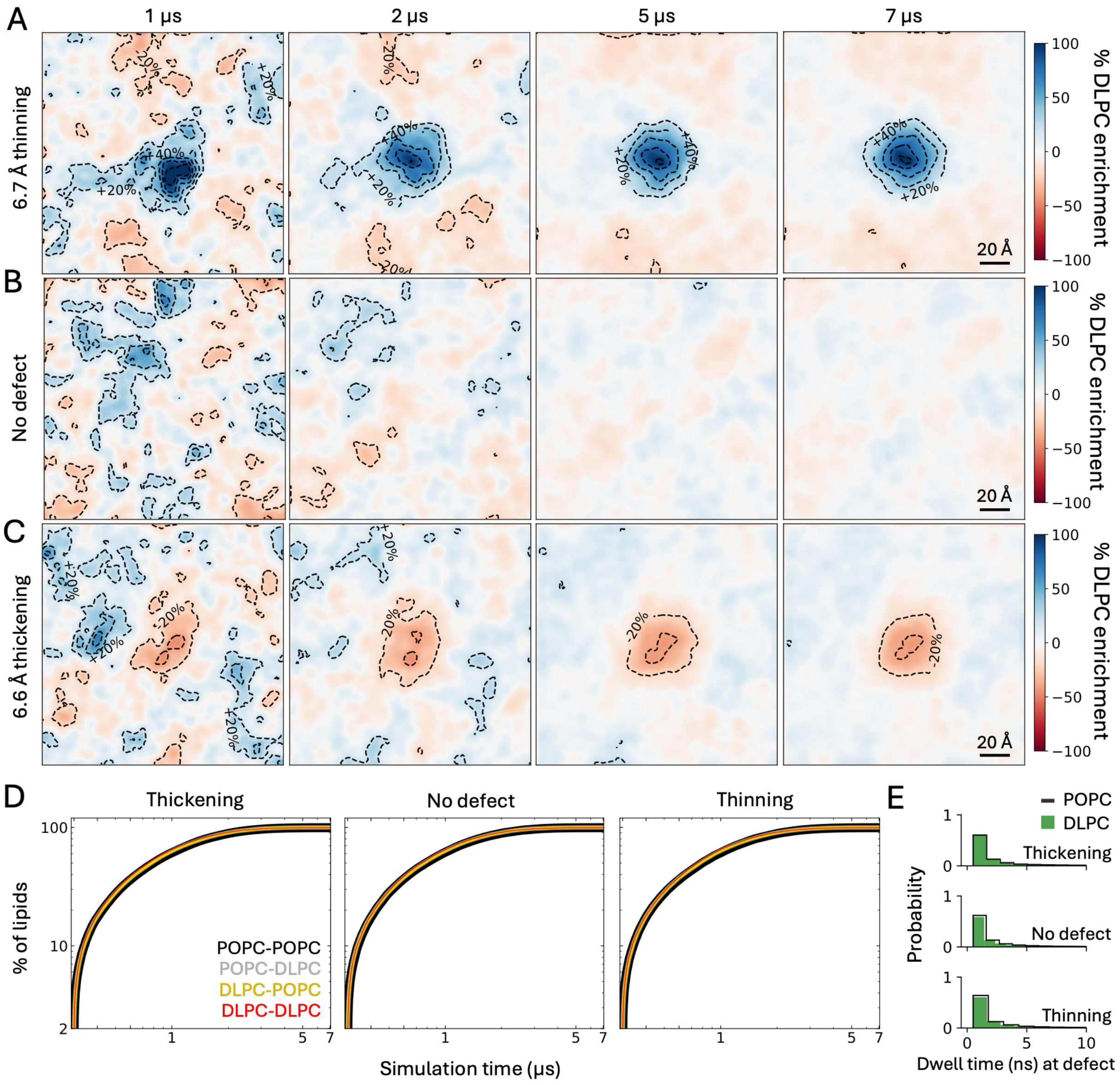


**Figure S7. Lipid mixing and statistical convergence in simulations of POPC:DLPC membranes**. (**A**) Distribution of DLPC lipids in a 1:1 POPC:DLPC membrane with a defect at its center that makes the membrane thinner, after 1, 2, 5, or 7 μs of simulation. The area of the defect is indicated. An enrichment level of 0% corresponds to the bulk ratio, i.e. 50%. Dashed contour lines denote enrichment increments of 20%. (**B**) Same as (A), without the thickness defect. (**C**) Same (A), when the defect induced in the membrane center makes the membrane thicker. (**D**) For each of the conditions represented in (A, B, C), the plots quantify the degree of lipid mixing, which we define as the percentage of lipids $j$ of a certain type (POPC or DLPC) that reside for at least 10 ns in a 12-Å shell around each lipid $i$ of the same or different type. Full mixing is reached when this quantity is 100% i.e. all the possible lipid pairs $ij$ in each leaflet have been observed to be first-neighbors for at least 10 ns. Note this condition is nearly achieved after 5 μs of simulation, irrespective of deformation magnitude. (E) Distribution of lipid dwell times in the thickness defect, in the same conditions represented in (A, B, C). Bin widths for POPC and DLPC are slightly different for clarity of visualization.
